# Sex differences in prenatal development of neural complexity in the human brain

**DOI:** 10.1101/2022.11.21.517302

**Authors:** Joel Frohlich, Julia Moser, Katrin Sippel, Pedro A. M. Mediano, Hubert Preissl, Alireza Gharabaghi

**Affiliations:** Institute for Neuromodulation and Neurotechnology, University Hospital and University of Tübingen, Germany; IDM/fMEG Center of the Helmholtz Center Munich at the University of Tübingen, University of Tübingen, German Center for Diabetes Research (DZD e.V.), Tübingen, Germany; Masonic Institute for the Developing Brain, University of Minnesota, Minneapolis, MN, USA; Department of Computing, Imperial College London, London, UK; Department of Psychology, University of Cambridge, Cambridge, UK; German Center for Mental Health (DZPG), Tübingen, Germany

**Keywords:** Fetus, Infant, MEG, Complexity, Development, Sex

## Abstract

The complexity of neural activity is a commonly used read-out of healthy functioning in cortical circuits. Prior work has linked neural complexity to the level of maternal care in preterm infants at risk for developing mental disorders, yet the evolution of neural complexity in early human development is largely unknown. We hypothesized that cortical dynamics would evolve to optimize information processing as birth approaches, thereby increasing the complexity of cortical activity. To test this hypothesis, we conducted the first ever study relating prenatal neural complexity to maturation. MEG recordings were obtained from a sample of fetuses and newborns, including longitudinal data before and after birth. Using cortical responses to auditory irregularities, we computed several entropy measures which reflect the complexity of the MEG signal. Despite our hypothesis, neural complexity significantly decreased with maturation in both fetuses and newborns. Furthermore, we found that complexity decreased significantly faster in male fetuses for most entropy measures. Our surprising results lay the groundwork for the first ever mapping of how neural complexity evolves in early human development, with important implications for future efforts to develop predictive biomarkers of psychiatric disorders based on the complexity of perinatal MEG signals.

## 1. Introduction

The entropy or “complexity” of neural signals has been proposed as a useful readout of cortical circuit functioning. Across a wide variety of clinical contexts, including neuropsychiatric disorders (1–3) and neurodegenerative disorders (4, 5), low entropy has been associated with, or even predictive of (1), impaired functioning. Similarly, in healthy individuals, low entropy is associated with states with a low capacity for information processing, such as non-rapid eye movement (NREM) sleep (6, 7) or general anesthesia (8, 9). On the other hand, high entropy is sometimes associated with abundant flexibility in circuits, e.g., in healthy individuals under the influence of psychedelic drugs (10, 11). A closely related measure of cortical dynamics, known as criticality, also follows a similar pattern from general anesthesia to the psychedelic state (12). The proper interpretation of neural entropy is open to some debate, yet these lines of evidence, combined with analogous findings from other organ systems (13, 14), suggest that optimal levels of physiological complexity might be general signatures of adaptability, flexibility, and/or efficient information processing, with potential for predicting mental health outcomes (1, 15).

In order to explore neural complexity in a developmental context, the meaning of neural signal entropy in prenatal and neonatal development should be better understood. Prior studies have examined preterm infants with this goal, given that preterm births are several times more likely to be associated with a later neuropsychiatric disorder (16–18). Two early electroencephalography (EEG) studies in infants found differences in signal complexity (measured as dimensionality) between sleep stages (suggesting differences in conscious level), as well as higher complexity in full-term infants compared with preterm infants who had reached the same postmenstrual age (PMA) (19, 20) (suggesting differences in cortical development). A later study reproduced this finding using different complexity measures and also found evidence that skin-to-skin care between mothers and preterm infants restores neural complexity in a small cohort (21). Next, another study that examined a similar intervention with a larger sample size (22) estimated EEG complexity in preterm infants as autoregressive integrated information (23). The authors (22) found that this measure relates to conscious state and increases with PMA in preterm infants during sleep; furthermore, this increase appeared greater for preterm infants who received more maternal care (22). While the results depended heavily on the choice of a time lag parameter used to estimate information integration, the overall finding appears to recapitulate an earlier study (24) that found increases with PMA in multiscale entropy of EEG signals recorded during sleep in preterm infants. More recently (25), the multiscale entropy of frontal EEG recorded from preterm infants during a mixture of spontaneous states (wakefulness and sleep) has also been shown to increase with PMA, while the multiscale entropy of functional near infrared spectroscopy (fNIRS) signals recorded from the same scalp site in the same infants decreased with PMA. Nonetheless, even in fNIRS data, complexity may also be clinically relevant: a previous study showed that high fNIRS signal complexity is associated with better acute outcomes (i.e., higher rates of survival) for very to extremely preterm infants in the neonatal intensive care unit (26).

Given the above evidence that skin-to-skin care influences neural complexity in preterm infants, a better understanding of how neural complexity evolves from late gestation to early infancy in healthy subjects may be useful for eventual efforts toward developing markers of treatment response in preterm infants at greater risk of developing mental disorders later in childhood (16). Although several of the above studies have already reported maturational changes in neural complexity in preterm infants, preterm infants are imperfect models of healthy development in age-equivalent fetuses, as the former inhabit a very different ex utero environment and are often forced to leave the womb early due to an in utero insult (27). Furthermore, while the above studies focused on spontaneous neural signals, investigations of perinatal neural complexity under conditions of controlled thalamocortical perturbation may be valuable for probing circuit development (28). Our current study addresses these research directions by examining neural entropy from event-related MEG signals in human fetuses and newborns, with the hypothesis that MEG signal complexity, like anatomical complexity (29), should increase with age both before and after birth. We measured cortical responses to auditory irregularities using 101 MEG recordings that passed rigorous quality control in a cohort with prior evidence of P300-like neural responses (30, 31), including 43 fetuses and 20 newborns; 16 subjects gave longitudinal data both before and after birth. Because sensory “oddballs” generate Bayesian prediction errors, we reasoned that they should more strongly perturb the thalamocortical system than repetitive trains of identical stimuli. Our approach thus takes inspiration from the perturbational complexity index (often referred to as PCI) (6, 8), which emphasizes causal influences within a system (32) [however, our method also features important differences, see (28)].

Surprisingly, we found evidence that neural entropy declines with maturation in both fetuses and newborns, and that this decline cannot be easily explained by noise differences according to a decision tree for classifying neural dynamics (33). The entropy decline occurs faster for male fetuses, corroborating earlier work suggesting sex differences in sensory evoked fetal MEG signals (34). The evolution of neural complexity in early development and its relationship to fetal sex should be understood prior to efforts toward using neural complexity as an in utero marker of prematurity and neuropsychiatric risk.

## Results

To map the evolving complexity of cortical dynamics, we estimated complexity using several distinct entropy measures, as well as decompositions of the MEG signal into amplitude and phase components and comparisons of the MEG signal entropy to that of surrogate signals. We therefore computed entropy from event-related data (28) and utilized auditory sequences with and without oddballs. Each experiment exposed fetuses and newborns to two blocks of stimuli during an exposure phase which followed different rules: sequences of identical tones (‘ssss’) or nonidentical tones (‘sssd’). In a subsequent test phase, sequences occasionally violated the block rule. For an overview of the experiment, see Figure 1. For additional details, see Methods and Materials.

**Figure 1:**
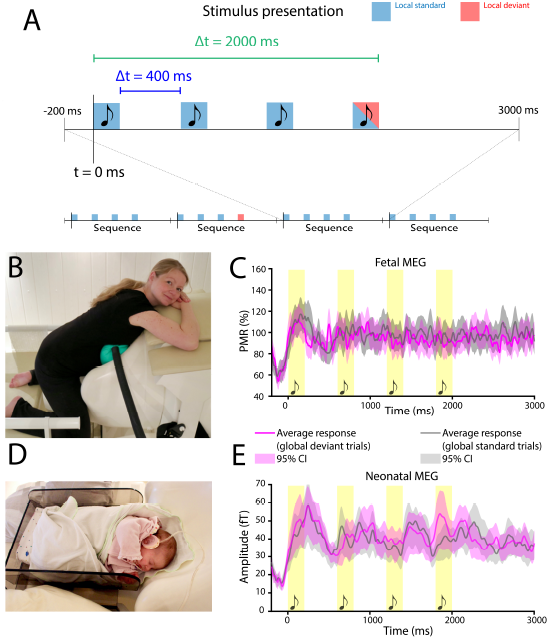
Overview of the experiment. Musical notes denote auditory stimuli. (A) Each auditory sequence consisted of four tones of 200 ms duration each, separated by a 400 ms intertone interval. The entire sequence, from the onset of the first tone to the offset of the fourth tone, was 2000 ms in duration. The fourth tone of each sequence varied during the test phase. After averaging across trials within each condition, we analyzed signals starting from 200 ms prior to the onset of the first tone to 1000 ms following the offset of the fourth tone (3200 ms duration total). (B) For fetal recordings, the expecting mother-to-be must position her abdomen within the concavity of the MEG the sensor array, with a sound balloon placed in between her body and the SARA device to deliver auditory tones. (C) Fetal MEG signals are recorded noninvasively in response to auditory tones. To correct for the influence of fetal head orientation and size on MEG signal amplitude, all signals were normalized as percent maximum response (PMR) relative to the maximum amplitude recorded during the earlier exposure phase. The average across all recordings for global and standard deviant trials are shown in magenta and gray, respectively. (D) After birth, a subset of subjects returned to the laboratory as newborns and were recorded from after being placed in a cradle oriented head-first toward the SARA device’s SQUID magnetometer array. In order to safely deliver auditory stimuli to the neonatal brain, the newborn wore infant-friendly headphones. (E) The SARA device records cortical signals noninvasively from newborns; again, the average of all global deviant trials is shown in magenta, and the average of all gloabl standard trials is shown in gray. Note that B and D are adapted from (28).

Specifically, we hypothesized that MEG entropy would increase with age in both fetuses (using gestational age) and newborns. Additional predictors such as stimulus sequence type (i.e, ‘ssss’ or ‘sssd’), block rule, heart rate variability (i.e., the standard deviation of heart beat intervals, SDNN, as an index of arousal), sex, PMA at birth, maternal age, and maternal body mass index (BMI) prior to pregnancy were also considered by a stepwise modeling procedure. Finally, to better understand the relationship between entropy and spectral properties, we correlated entropy with spectral power and the spectral exponent, and we also analyzed event related spectral perturbations (ERSPs).

We utilized data that were collected from a previous study at the fMEG Center at the University of Tübingen. Recorded signals included both cortical activity (MEG, bandpass filtered 1 - 10 Hz in fetuses and 1 - 15 Hz in newborns) and cardiac activity (magnetocardiography or MCG), the latter of which was retained separately to measure HRV [one of the main parameters used to classify fetal behavioral and sleep states (35)] and, by proxy, arousal. The dataset consisted of 81 usable recordings of cortical and cardiac signals from 43 fetuses (gestational age range: 25 – 40 weeks) which passed strict MEG quality control (74.3% of recordings retained). To correct for the confounding influence of differences in fetal head size and orientation, we normalized fetal MEG signals as percent maximal response (PMR) relative to MEG recorded during the earlier exposure phase of the experiment (Fig. 1). Based on previous simulations, we estimate that the SNR of fetal MEG signals after preprocessing and trial-averaging was approximately 10 dB (i.e., 3:1) (36) and much higher for neonatal MEG signals. Of the data which survived quality control, 75 recordings from 41 fetuses also contained usable MCG data, allowing us to estimate HRV; however, since HRV was not selected a predictor of entropy by a stepwise modeling procedure, all 81 recordings from 43 fetal subjects were used in fetal entropy models regardless of whether they contained usable MCG.

We additionally included 20 recordings from 20 newborns (age range: 13 – 59 days) acquired with the same MEG system and which also passed strict MEG quality control (60.1% of recordings retained); 16 newborns were also recorded from prior to birth in the fetal cohort. All newborns in our sample had full-term births.

We estimated the complexity of MEG signals averaged across trials and channels according to six different entropy measures (see Materials and Methods; note that channel-averaging is not principally motivated by low SNR but, rather, by variable head positions between recordings, which precludes direct comparisons of data from a specific channel between recordings, even within the same subject, as the fetal or neonatal head may have been oriented differently in each instance. Two entropy approaches, Lempel-Ziv complexity (LZC) (37) and context tree weighting (CTW) (38), measure the compressibility of the signal. The remaining approaches are based on state-space reconstruction of the signal and include modified sample entropy (mSampEn, or the tendency of motifs to reoccur within a signal) (39), modified multiscale entropy (mMSE, which computes mSampEn at different time scales) (40), and permutation entropy (PermEn, or the occurrence of unique permutations based on ordinal rankings of data) (41). We computed two different varieties of PermEn using a 32 ms lag (PermEn32, sensitive to 4 - 10 Hz activity) and a 64 ms lag (PermEn64, sensitive to 2 – 5 Hz activity).

For each entropy measure, we selected independent variables using stepwise regression and then built a common model for all entropy measures (done separately for fetuses and newborns), entering features into the common model if they were selected by the stepwise procedure for at least half of the entropy features; the common model also used random intercepts to account for repeated measures in a linear mixed model (LMM) framework. Because fetal and neonatal data were filtered and preprocessed differently, primary analyses were performed on each separately; however, for later analyses in which data from both fetuses and newborns were pooled together (including longitudinal measurements), we applied a second filter to neonatal recordings to match the 10 Hz lowpass frequency cutoff of fetal recordings. This also allowed us to visualize entropy trajectories for both fetal and neonatal data as a function of PMA ranging from pre- to post-birth (Fig. 2). Finally, we also used a stepwise procedure to model power at each time-frequency element of ERSPs and corrected for multiple testing across time-frequency elements using threshold-free cluster enhancement (42). For all P-values generated by LMMs outside of ERSPs, we used the Benjamini-Hochberg (BH) procedure to control the false discovery rate (FDR) at 0.05, resulting in *P*_*crit*_ = 0.024. Other statistical tests (e.g., simple correlations, chi-squared tests, etc.) were either uncorrected or P-values were not computed, as these tests were not the main focus of our analysis.

**Figure 2:**
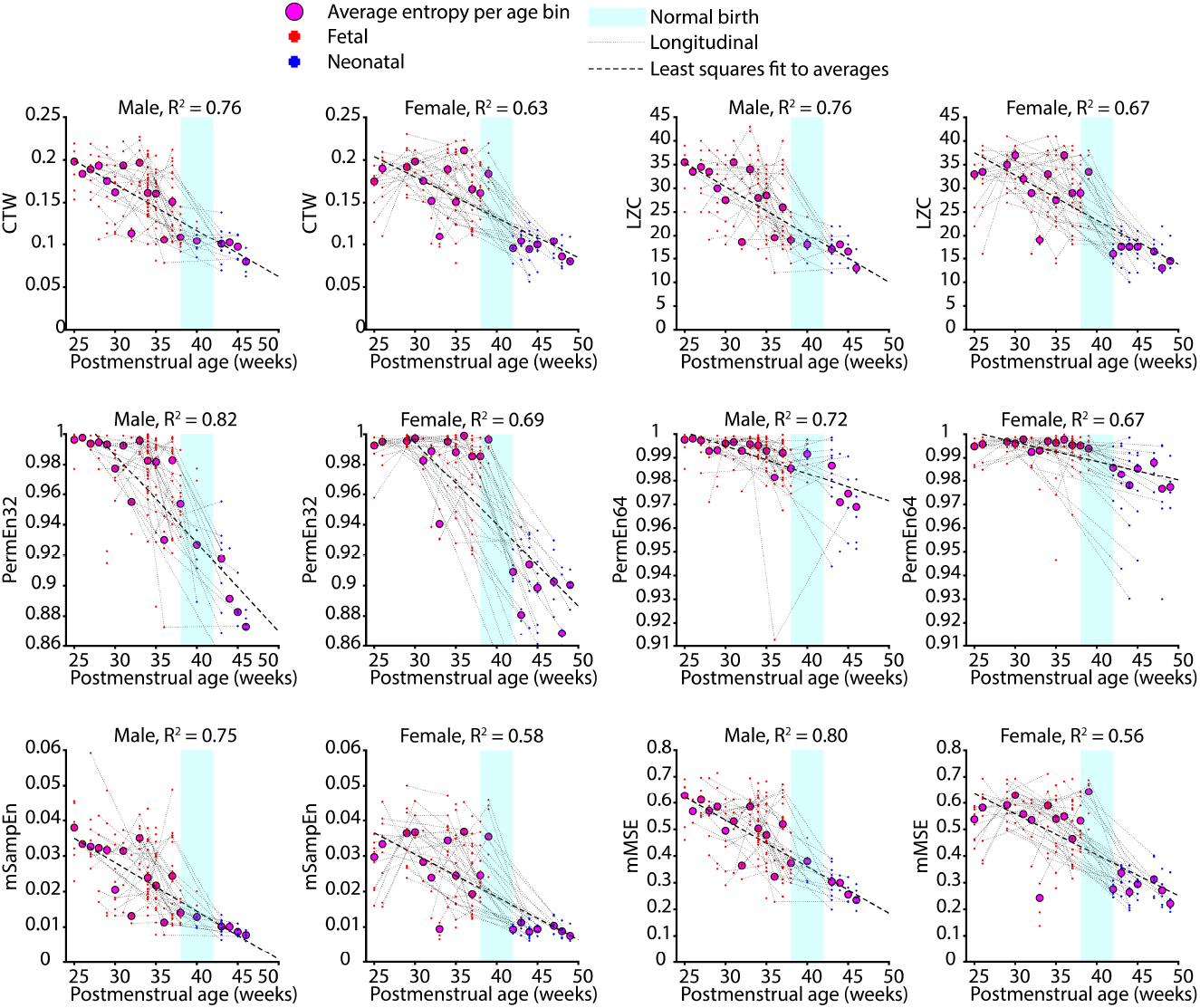
Scatter plots of postmenstrual age versus signal entropy. Red data points (fetuses) and blue data points (newborns) are taken from each session and rule/stimulus condition, with thin dotted lines connecting longitudinal data from the same subjects. All neonatal data were filtered at the same frequency as fetal data such that neonatal signal entropy estimates could be compared with fetal entropy estimates in the same space, revealing a continuous entropy decline with postmenstrual age. Time of normal birth is indicated in cyan. We averaged data both within- and between-subjects at the level of one-week time bins (magenta circles) and computed the variance explained (*R*^2^) according to the least-squares fit of the smoothed data. Because of the GA x sex interaction found in fetal data, we plotted data separately for males (first and third columns) and females (second and fourth columns). The above data show that signal entropy follows a smooth, continuous decline ranging from gestation to after birth. In all cases, postmenstrual age explains the majority of variance in signal entropy.

### MEG entropy declines with maturation in fetuses and newborns, with a faster decline in male fetuses

In fetuses, the stepwise procedure used to build LMMs selected the following fixed effects: gestational age (GA), maternal age (MA), sex, and the interaction of sex and GA (note: here, we use the term ‘GA’ to refer to age since last menstruation only in fetuses, whereas in cases where neonatal data are combined with fetal data, we use the term ‘PMA’ to avoid confusion with the subject’s gestational age at birth). All six entropy measures were significantly predicted by GA (*P <* 0.001), and in all cases, entropy *declined* with increasing GA, contrary to our hypothesis. Several entropy measures (CTW, LZC, mMSE, and mSampEn) featured a significant sex by GA interaction (*P < P*_*crit*_), with a greater entropy decline in males, and both PermEn measures featured trend-level interactions in this direction (*P <* 0.05) which failed to survive the BH procedure for controlling the FDR. In half of our entropy measures (CTW, mSampEn, and mMSE), we observed a main effect of fetal sex (significantly lower entropy in male fetuses, *P < P*_*crit*_), with a similar trend-level effect for LZC and PermEn64 (*P <* 0.05). MA did not significantly predict entropy as quantified by any measure. See Table 1 for exact P-values.

**Table 1.** Fetal entropy models. GA = gestational age, MA = maternal age.

| Measure | CTW | LZC | PermEn32 | PermEn64 | MSE | SE |
| --- | --- | --- | --- | --- | --- | --- |
| GA P-value | 1.05e-06 | 5.47e-06 | 4.99e-06 | 0.000931 | 1.44e-05 | 8.55e-07 |
| GA t-stat | -5.01 | -4.65 | -4.67 | -3.35 | -4.43 | -5.06 |
| GA survive FDR | TRUE | TRUE | TRUE | TRUE | TRUE | TRUE |
| MA P-value | 0.152 | 0.323 | 0.467 | 0.128 | 0.0946 | 0.0923 |
| MA t-stat | 1.44 | 0.991 | 0.729 | 1.53 | 1.68 | 1.69 |
| MA survive FDR | FALSE | FALSE | FALSE | FALSE | FALSE | FALSE |
| Sex x GA P-value | 0.00278 | 0.0129 | 0.0678 | 0.0289 | 0.00572 | 0.00534 |
| Sex x GA t-stat | 3.02 | 2.5 | 1.83 | 2.2 | 2.79 | 2.81 |
| Sex x GA survive FDR | TRUE | TRUE | FALSE | FALSE | TRUE | TRUE |
| Sex P-value | 0.00798 | 0.0299 | 0.162 | 0.0488 | 0.0122 | 0.0135 |
| Sex t-stat | -2.68 | -2.19 | -1.4 | -1.98 | -2.53 | -2.49 |
| Sex survive FDR | TRUE | FALSE | FALSE | FALSE | TRUE | TRUE |

In newborns, the stepwise procedure selected only age as a fixed effect predictor of entropy. For half of the entropy measures we examined (PermEn32, mSampEn, CTW), entropy significantly decreased with age in newborns (*P < P*_*crit*_), and trends were also present for LZC and mMSE (*P <* 0.05). For exact P-values, see Table 2. Because each subject was only recorded from once as a newborn, the above relationship between age and entropy in early infancy is entirely correlational and cannot be verified using longitudinal data; however, it matches with our finding of declining entropy with GA in fetuses (see above), which is based on both correlational *and* longitudinal data.

**Table 2.** Neonatal entropy models.

| Measure | CTW | LZC | PermEn32 | PermEn64 | MSE | SE |
| --- | --- | --- | --- | --- | --- | --- |
| age P-value | 0.023 | 0.0383 | 0.00162 | 0.146 | 0.0434 | 0.0119 |
| age t-stat | -2.33 | -2.11 | -3.28 | -1.47 | -2.06 | -2.58 |
| age survive FDR | TRUE | FALSE | TRUE | FALSE | FALSE | TRUE |

Next, we considered the entropy of neonatal MEG signals filtered at the same frequency as fetal signals, which allowed us to combine both fetal and neonatal data on a continuum of PMA. Entropy measures were averaged both between- and within-subjects in one-week age bins, and after this data smoothing, the change in entropy with PMA was considered separately for male and female subjects. In all cases, PMA explained the majority of variance in entropy (*R*^2^ *>* 0.5), and for each entropy measure, the *R*^2^ was larger for male subjects (Figure 2). The largest proportion of entropy explain by PMA was PermEn32 in males (*R*^2^ = 0.82).

To assess the extent to which measurements within-subjects were still similar to each other when data were pooled across fetuses and neonates, we measured intraclass correlations (ICC) for each entropy measure, performed separately for pooled data and fetal data alone. In nearly all cases, ICCs demonstrated that entropy measurements were fairly consistent (*ICC >* 0.4) within subjects across time, except for PermEn64, which yielded poor consistency (*ICC <* 0.3). Crucially, in all entropy measures except for PermEn64, the ICC did not decrease when data were pooled across fetuses and newborns, but rather increased slightly (Table 3), suggesting that measurements from pre- and post-birth can be considered on the same continuum of development.

**Table 3.** Intraclass correlations (ICCs) with P-values computed from nonparametric tests with 1000 permutations. Including neonatal data together with fetal data was only detrimental to the ICC corresponding to PermEn64.

| Measure | ICC (fetal only) | P-value (fetal only) | ICC | P-value |
| --- | --- | --- | --- | --- |
| CTW | 0.441 | < 0.001 | 0.446 | < 0.001 |
| LZC | 0.435 | < 0.001 | 0.436 | < 0.001 |
| PermEn32 | 0.478 | < 0.001 | 0.490 | < 0.001 |
| PermEn64 | 0.269 | 0.002 | 0.255 | 0.031 |
| MSE | 0.478 | < 0.001 | 0.520 | < 0.001 |
| SE | 0.455 | < 0.001 | 0.483 | < 0.001 |

Finally, we considered the pooled fetal and neonatal data with entropy modeled using LMMs. The stepwise procedure selected fixed effects of PMA and sex, in addition to a categorical variable BIRTH included in the default model indicating whether data were recorded from fetuses or newborns. All six entropy measures significantly declined with PMA (*P <* 0.0001) and were smaller for newborns than for fetuses (*P <* 0.01). Although sex was entered into the model as a fixed effect, it did not significantly predict entropy for the pooled data (but note a trend of lower PermEn32 in males, *P = 0*.*03*). For exact P-values, see Table 4.

**Table 4.** Entropy models (fetal and neonatal data combined). Birth is a categorical variable denoting whether subjects are born yet (NO = fetuses; YES = newborns). PMA = postmenstrual age.

| Measure | CTW | LZC | PermEn32 | PermEn64 | mMSE | mSampEn |
| --- | --- | --- | --- | --- | --- | --- |
| Birth P-value | 4.93e-09 | 6.38e-09 | 0 | 8.82e-05 | 2.94e-11 | 8.81e-07 |
| Birth t-stat | -6.02 | -5.97 | -11.8 | -3.97 | -6.9 | -5.02 |
| Birth survive FDR | TRUE | TRUE | TRUE | TRUE | TRUE | TRUE |
| PMA P-value | 6.14e-07 | 7.24e-07 | 2.77e-07 | 0.00561 | 1.53e-05 | 2.38e-07 |
| PMA t-stat | -5.09 | -5.06 | -5.25 | -2.79 | -4.39 | -5.28 |
| PMA survive FDR | TRUE | TRUE | TRUE | TRUE | TRUE | TRUE |
| Sex P-value | 0.0529 | 0.0779 | 0.0298 | 0.08 | 0.259 | 0.127 |
| Sex t-stat | 1.94 | 1.77 | 2.18 | 1.76 | 1.13 | 1.53 |
| Sex survive FDR | FALSE | FALSE | FALSE | FALSE | FALSE | FALSE |

### Changes in MEG signal amplitude drive maturational decreases in signal entropy

To determine whether the decline in entropy with maturation was driven by changes in the MEG signal amplitude or, alternatively, by non-amplitude changes (i.e., changes in the MEG signal phase and its interaction with amplitude), we performed an entropy decomposition (43), which entailed randomizing Fourier phases to isolate the contributions of amplitude, phase, and their interaction (see Materials and Methods). In fetal MEG recordings, we performed this analysis within-subjects using 12 fetuses with longitudinal data from the lower and upper one-third of the GA distribution. In neonatal MEG recordings, we performed this analysis between 10 younger subjects in the lower half of the age distribution and 10 older subjects in the upper half of the age distribution.

In both fetuses and newborns, we found that, for nearly all entropy measures, changes in the amplitude properties of the signal diminished entropy significantly more so than changes in non-amplitude properties (Table 5, *P < P*_*crit*_), i.e., the MEG signal amplitude was responsible for the maturation-related decline in entropy. In fact, using PermEn32 and PermEn64, we found an extremely significant effect (*P <* 10^−32^) whereby changes in non-amplitude properties of the signal always resulted in increases in PermEn, whereas changes in amplitude properties of the signal always resulted in decreases in PermEn. The huge magnitude of this effect (Cohen’s d > 20 in fetuses and newborns for both PermEn32 and PermEn64, see Fig. S1) is likely due to PermEn’s unique sensitivity to ordinal rankings of data, meaning that very small signal changes should affect PermEn to a greater degree than the other entropy measures we employed. Perhaps for this reason, LZC in fetuses and CTW in newborns did not yield significant differences between amplitude and non-amplitude signal components (but note a trend-level effect for neonatal CTW, *P <* 0.05).

**Table 5.** Entropy decomposition. LR stat = log-likelihood ratio test statistic.

| Measures | CTW | LZC | PermEn32 | PermEn64 | mMSE | mSampEn |
| --- | --- | --- | --- | --- | --- | --- |
| Fetal P-value | 0.0242 | 0.0548 | < 1e-32 | <1e-32 | 0.00387 | 0.00984 |
| Fetal LR stat | 5.08 | 3.69 | 271 | 258 | 8.34 | 6.66 |
| Fetal survive FDR | TRUE | FALSE | TRUE | TRUE | TRUE | TRUE |
| Neonatal P-value | 0.0271 | 0.0147 | <1e-32 | <1e-32 | 0.0207 | 0.0174 |
| Neonatal LR stat | 4.89 | 5.95 | 369 | 405 | 5.35 | 5.66 |
| Neonatal survive FDR | FALSE | TRUE | TRUE | TRUE | TRUE | TRUE |

### MEG amplitude does not significantly mediate the effect of fetal sex on signal entropy

Given the results of the entropy decomposition, we next asked if signal amplitude mediated the effect of fetal sex on entropy. Using a nonparametric mediation analysis (44), we treated sex as the independent variable, the mean signal amplitude as the mediator variable, and each of our six entropy measures as the dependent variable in separate models fit to fetal data. In all cases, we found no significant effect of mediation, even without accounting for multiple testing (see Table S1).

### Evidence of cortical stochasticity in fetuses and neonates

To assess the degree to which cortical MEG signals differed from noise, i.e., surrogate signals with the same amplitude distribution, we next tested whether the entropy estimates for cortical signals were significantly different from the entropy of surrogates. In each case, we modeled entropy separately in fetuses and newborns using surrogacy as a predictor in LMMs alongside the same fixed and random effects as were used in the previous entropy models. Surrogacy did not significantly predict any entropy measure except for mSampEn in newborns (*P < P*_*crit*_, higher entropy in surrogate signals; see Table S2 for exact P-values); note also a trend-level effect for PermEn32 in fetuses (*P <* 0.05, lower entropy in surrogate signals). This generally null result suggests that MEG signals lack nonlinear components that affect entropy.

Given this likely lack of nonlinearity, at least some degree of stochasticity, or intrinsic randomness, appears to be present in MEG signals. Using the Toker decision tree algorithm (33), we found that the proportion of signals with deterministic dynamics was significantly larger in fetuses than in newborns (i.e., neonatal signals were more likely to show intrinsically random behavior; *χ*2 = 28.6, *P* = 9.1 · 10^−8^); note that each recording was treated as an independent sample). When we applied the same predictor from Table 1 and Table 2 to model signal dynamics in fetuses and newborns, respectively, no predictor showed a significant effect, suggesting that our finding of declining entropy with maturation cannot be easily explained by random noise. While we did observe a trend-level effect of more stochastic dynamics in older fetuses (P = 0.06) consistent with our finding that newborns are more likely to exhibit stochastic dynamics than fetuses, this relationship is rather the opposite of what one should expect if the age-related decline in entropy is driven by declining noise (see Discussion).

### Entropy measures correlate strongly with one another and weakly with spectral power

To better understand the behavior of signal entropy measures in our data, we computed correlations between these measures. Because some fetuses gave data at multiple laboratory visits, we first evaluated whether these recordings could be approximated as independent samples by examining the usefulness of the random intercept term in LMMs. For each fetal model except for the LMM predicting PermEn64, the random effect term significantly increased the model fit (*P <* 10^−6^, log-likelihood ratio test, Table S4), implying substantial dependencies between longitudinal recordings from the same subjects. This result is in agreement with ICCs showing consistency within longitudinal recordings from the same subjects (Table 3). For this reason, fetal correlations were estimated using the beta coefficeints of LMMs with random intercepts (see *Materials and Methods*).

As expected, in both fetal and neonatal data, entropy measures were positively correlated with one another (Fig. S3A,B), many strongly so (fetal range: *β* = 0.42 – 0.97; neonatal range: r = 0.61 – 0.97). This shows that these measures are approximately capturing a reliable underlying property of the MEG signal. Although we used beta coefficients to measure correlations in fetal data with longitudinal recordings, beta coefficients returned very similar values as Pearson coefficient (Fig. S4). Next, we examined correlations between entropy measures and spectral power. Entropy measures were negatively correlated with spectral power at all frequencies in newborns (range: r = -0.77 – -0.089) and at frequencies < 5.5 Hz in fetuses (range: *β* =-0.48 – -0.09), suggesting that cortical synchronization introduces regularities into cortical signals that constrain their entropy (Fig. S3C,D). However, this relationship between entropy and power reversed at faster frequencies in fetuses, with moderate correlations as high as *β* = 0.36 for power at 9.5 Hz.

Next, we utilized time-resolved CTW to evaluate the relationship between entropy and spectral power within each recording. This within-recording analysis was only performed using CTW, given its high temporal resolution (45); we accepted this analysis as being representative of entropy in general, given that CTW is highly correlated with other entropy measures (Fig. S3, S4). We found that, after averaging timefrequency representations (TFRs) and time-resolved CTW across all datasets, CTW was not strongly correlated with spectral power at most frequencies in fetuses and newborns; note that data were averaged in a nested fashion (first with-subjects, then between-subjects) to ensure that subjects with longitudinal data were not overrepresented. After averaging correlation coefficients across all four stimulus x block rule combinations, we found that the absolute value of the mean correlation did not exceed 0.25 for any frequency in fetal or neonatal data (Fig. S3E,F). Thus, signal entropy appears to be weakly correlated with spectral power within each dataset, but negatively—and, in some cases, strongly—correlated with spectral power between datasets (Fig. S3).

Having examined correlations between entropy and spectral power, we next measured the slope (i.e., spectral exponent) of the aperiodic component of MEG signals and correlated the spectral exponent with entropy estimates in both newborns and fetuses (Fig. S3G). The spectral exponent is a useful approximation of the balance between excitatory and inhibitory synaptic transmission (46), and the spectral exponent is known to increase in states of reduced arousal and consciousness (47). In our data, correlations between the spectral exponent and entropy estimates were negative, ranging from r = -0.49 (correlation with PermEn64 in fetuses) to r = -0.83 (correlation with mSampEn in newborns). Given that MEG signal entropy is negatively correlated with both maturation (Fig. 2) and the spectral exponent (Fig. S3G), the relationship between the former and the latter is positive; the spectral exponent is weakly correlated with GA (r = 0.20) and age (r = 0.34) in fetuses and newborns, respectively. However, given that MEG signals were lowpass filtered at a low cutoff frequency of 10 Hz in fetuses and 15 Hz in newborns, we did not have enough bandwidth to fit the spectral exponent to the slope of both low frequency and high frequency activity, as is often done in other studies (48, 49). Thus, given our bandwidth limitation, we opted not to rigorously model and test relationships between the spectral exponent and developmental variables such as GA, age, or sex.

### Cortical synchronization increases with maturation in fetuses and newborns

Finally, we examined how developmental variables influenced ERSPs in fetal and neonatal data. Using lasso regression, we identified predictor variables to enter into LMMs as fixed effect predictors of spectral power. We then applied threshold-free cluster enhancement (TFCE) to derive empirical P-values corrected for the large number of time-frequency elements that each fixed effect was tested across. Results from ERSPs revealed a broad increase in neural synchronization with maturation in both fetuses and newborns. In fetal data (Fig. 3A), we observed a significant increase (*P*_*TFCE*_ *<* 0.01) in low frequency (< 5 Hz) synchronization with GA at most timepoints; conversely, we also observed several clusters of diminished spectral power with GA for relatively high frequencies (> 6 Hz) at many timepoints. Fetal data also showed significant (*P*_*TFCE*_ *<* 0.01) synchronization in the canonical theta band [4 - 8 Hz, more likely an alpha rhythm in the perinatal brain (50)] related to local deviant tones approxi- mately 400 - 750 ms after the onset of the fourth tone (Fig. 3B). On the other hand, we also found what would be, if taken at face value, a desynchronization effect of stimulus prior to the fourth tone, which may be better understood as an artifact of the normalization process applied to fetal signals (i.e., signals with large responses after the fourth tone were shifted down in amplitude by the normalization, causing an artifactual decrease in amplitude before the fourth tone).

**Figure 3:**
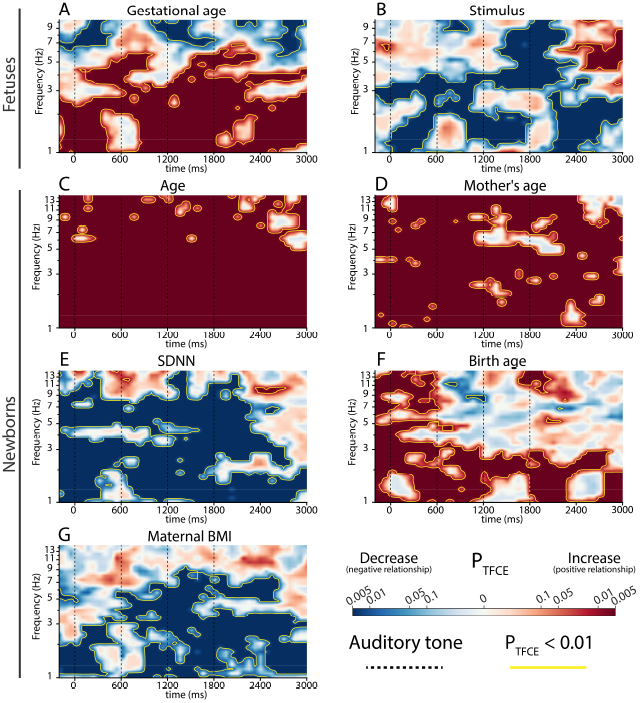
Effects of developmental variables and stimuli on ERSPs. P-values from threshold-free cluster enhancement (*P*_*TFCE*_) are color coded according to size (darker colors represent smaller P-values) and the sign of the effect (red colors represent positive relationships with spectral power, blue colors represent negative relationships with spectral power). Yellow contours denote clusters that statistically significant at *P*_*TFCE*_ < 0.01. (A) GA generally corresponds to low-frequency synchronization and high-frequency desynchroization in fetuses. (B) Local deviant trials (fetuses) result in 4 - 8 Hz synchronization in fetuses beginning 400 - 750 ms after the onset of the final tone and persisting until the end of the trial. Also note that our finding of desynchornization prior to the final tone is a normalization artifact (recall that we normalized fetal signals according to the maximum amplitude observed during the exposure phase). (C) In newborns, synchronization significantly increased for virtually all frequencies and timepoints with age. (D) Unexpectedly, MA also related to synchronization in newborns across nearly all frequencies and timepoints. (E) SDNN (a proxy for arousal) corresponded to MEG desynchronization at many frequencies (especially delta) and timepoints in newborns, as would be expected if low SDNN denoted a sleep state marked by cortical synchronization and delta activity. (F) ERSPs were predicted by PMA at which infants were borns; here, the developmental effects are similar to those of GA in fetuses. (G) The mother’s BMI (measured before pregnancy) corresponded to lower MEG synchronization, particularly at lower frequencies and timepoints later in the tone sequence.

In neonatal data, both the age of the newborn (Fig. 3C) and the age of the mother (Fig. 3D) significantly (*P*_*TFCE*_ *<* 0.01) predicted increased spectral power at nearly all frequencies and timepoints. Similarly, the newborn’s PMA at birth (Fig. 3F) was also significantly (*P*_*TFCE*_ *<* 0.01) related to spectral power: a later birth was predictive of increased power at nearly all frequencies prior to the second tone, and at most delta (< 3 Hz) frequencies thereafter.

The remaining two variables, neonatal SDNN and maternal BMI, both showed a significant (*P*_*TFCE*_ *<* 0.01) inverse relationship with cortical synchronization in newborns. Our finding in SDNN (Fig. 3E) is consistent with work showing that SDNN increases during states of cortical arousal in overnight sleep studies from adults (51) and the fact that cortical arousal should generally coincides with desynchronization mediated by acetylcholine (52). Finally, given that high maternal BMI is a known pregnancy risk factor (53–55), our finding of weaker cortical synchronization with increasing maternal BMI (which was measured prior to pregnancy) could be viewed as evidence that infants born to mothers with higher relative obesity exhibit less mature cortical activity.

In short, results from ERSPs demonstrate that an increase in spectral power (particularly at low MEG frequencies) with maturation generally coincides with the decrease in entropy we found earlier in our data, thus supporting the results of our entropy decomposition which showed that maturation-related decreases in signal entropy are driven by changes in signal amplitude (however, see *“Developmental effects on neural synchrony”* in the Discussion for a relevant caveat).

## Discussion

Herein, we built on previous studies which used electromagnetic perturbations in adults (6, 8) to probe the information processing capacity or “causal structure” (32) of neural circuits. Specifically, we used sensory perturbations in fetuses and newborns toward a similar purpose, as we first proposed in a recent manuscript (28). Although we built on an earlier proof-of-concept study (56), our current study is the first investigation to ever relate the entropy of human fetal brain dynamics to developmental variables such as (gestational) age or sex.

Surprisingly, and in contrast to studies of spontaneous neural entropy in EEG from preterm infants (24, 25), we found strong evidence that neural complexity declines with maturation in fetuses and neonates (with fullterm birth) in the context of sensory perturbations that generate prediction errors (30, 31), and that this decline happens more rapidly in male fetuses. Importantly, our findings cannot be easily attributed to decreasing noise with maturation because maturation did not significantly predict signal dynamics (stochastic versus deterministic) in fetuses or neonates (Table S2), suggesting that the level of stochasticity (i.e., intrinsic noise) does not change with maturation within either group (recall that entropy decreased both within and between groups). Furthermore, even when we consider the between group change in MEG entropy, which decreased after birth (Table 4), MEG signals from newborns contained a higher level of intrinsic noise (as inferred by stochastic dynamics, see Table S3, Fig. S2), which would otherwise imply higher entropy.

Although we studied a healthy sample of fetuses and newborns at normal risk for developing psychiatric disorders later in life, our entropy results may nonetheless be of interest for efforts toward developing predictive biomarkers of psychiatric disorders in early development for the following reasons 1) evidence from earlier studies has linked the complexity of neural activity to adaptability and health (1–3, 15, 57), 2) earlier studies show that preterm infants, who are at a higher risk of developing a psychiatric disorder (16–18, 58), show within-group differences in neural complexity based on level of maternal care (21, 22) and between-group differences in neural complexity compared with full-term infants (19, 20), and 3) female fetuses are generally less vulnerable to an adverse in utero environment (59), thus underscoring the relevance of our prenatal sex finding for predicting psychiatric disorders. Additionally, the spectral exponent of spontaneous neural activity during sleep has been reported to predict later autism risk in preterm infants (49), and in our data, MEG entropy was inversely related to the spectral exponent as estimated using narrow bandwidth spectra (Fig. S3G), thus suggesting a further avenue by which our results may facilitate the development of early predictive biomarkers.

### In utero sex differences in human brain activity

A small number of prior studies have reported in utero sex differences in human brain activity. Perhaps most relevant to our work, an investigation by another fetal MEG site (34) of auditory and visual evoked responses in MEG signals recorded from high-risk fetuses found that visual evoked responses (generated by shining red LED lights through the maternal abdomen) were detected significantly more often in male fetuses, with a similar trend for auditory evoked responses. These results are consistent with our finding of lower entropy in several measures of neural complexity in male fetuses (Table 1), as lower entropy signals should show a more pronounced evoked response which is easier to detect than responses in high-entropy waveforms, whose complexity may mask an evoked response (Fig. 4A).

**Figure 4:**
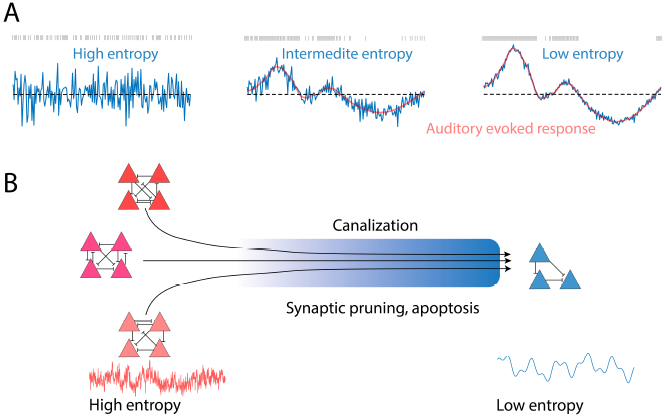
Possible scenarios explaining the decline of MEG entropy with perinatal maturation. (A) MEG signal entropy may decline with age in fetuses and newborns due to the maturation of auditory evoked responses, which should impose structure on the cortical signal, thus minimizing its entropy. Specifically, this scenario may explain decreases in entropy driven by amplitude changes with maturation. (B) The decline in neural complexity with maturation may reflect developmental processes such as apoptosis and synaptic pruning that reduce the number of ways in which neural circuits can be arranged (i.e., entropy). In this illustration, multiple circuit configurations (left) are possibly prior to the canalization process which brings circuits toward a genetically determined phenotype (right).

Besides MEG, functional magnetic resonance imaging (fMRI) has also been used to detect prenatal sex differences in human brain activity. As in our neural entropy data, Wheelock et al. (60) reported interactions between sex and GA in fMRI connecitivity network data from a large sample of human fetuses (25 to 39 weeks gestation) which “confirm that sexual dimorphism in functional brain systems emerges during human gestation.” A later study by Cook et al. (61) examined nearly 100 fetuses (19 to 40 weeks gestation) and used machine learning on fMRI connectivity data to classify fetal sex with 73% accuracy. Nonetheless, it remains unclear if and how these prenatal sexual dimorphisms in neural activity, including the one found in our study, relate to greater male vulnerability (59) to early developmental risk factors for later neuropsychiatric disorders (see *“Potential clinical impact”* below).

### Evidence of two opposing developmental processes

An entropy decomposition revealed that maturational changes in entropy are specifically driven by changes in signal amplitude with age, as best demonstrated using PermEn, which, unlike other entropy measures, is sensitive to differences in the ordinal rankings of data points. Furthermore, PermEn measures also show that non-amplitude properties of the MEG signal drive *increases* in entropy with maturation. Thus, the total observed effect of decreasing entropy with perinatal maturation may actually be the net result of two separate maturational processes, one which sculpts structured and predictable cortical responses to stimuli via increases in signal amplitude, thus lowering entropy, and another process which reduces the predictability of cortical responses via changes in signal phase, thus increasing entropy. The first process is consistent with results from ERSPs showing enhanced neural synchrony with maturation (Figure 3) and may plausibly result from forces which shape synaptic development and constrain neural circuit formation. The second process, however, remains elusive, though previous work in young children demonstrates that phasic EEG changes which affect signal entropy are related to transitions between wakefulness and NREM sleep (7); more generally, changes in EEG signal entropy often relate to one’s conscious state (62), thus suggesting that an increasing capacity for consciousness in developing brain networks drives the increase in neural entropy due to phasic MEG changes.

The maturation of ERF components, which introduce structure into cortical signals, thereby constraining their entropy (Figure 4A), may mask the effects of phasic changes on MEG signal entropy, a view supported by the fact that signal amplitude is responsible for the decrease in entropy (Fig. S1). Previous work in the same sample of fetuses which we studied here has shown maturation of P300-like cortical responses to deviant tones late in the third trimester, with fetuses demonstrating stronger responses past 35 weeks gestation as compared to younger fetuses (31). However, in later development (one month to adulthood), separate studies have shown that the entropy of event-related cortical signals increases as the evoked responses mature (63–65). The reason for this switch from decreasing to increasing neural entropy with maturation in the first year of life is unclear. One possible reason, synaptic pruning, is an important developmental event that only starts in the first months of life (66), yet we should expect pruning to further constrain the entropy of neural circuit activity (Fig. 4B). Alternatively, it is possible that different tasks and stimuli yield different entropy changes across development, some featuring increasing entropy and others featuring decreasing entropy. To the best of our knowledge, neural entropy recorded during a local-global paradigm has never been related to maturation in other age groups. While one recent machine learning study in older infants (57) did examine the entropy of auditory responses to linguistic oddball stimuli recorded with EEG (a similar, though not identical, task to that which we employed), the study focused on differences between diagnostic groups and did not report the relationship between entropy and age.

Our finding of a net decline in complexity with maturation was robust across six different entropy measures (Table 1), demonstrating that it generalizes across different estimators of complexity, albeit ones that are limited to time-domain signals. Although we were unable to investigate spatiotemporal patterns in our data due to variability in the fetal and neonatal head position between recordings, a previous study by Isler et al. (22) found changes in autogressive integrated information (a measure which accounts for spatial information) with maturation in preterm infants. We advocate for future studies which may use either MEG with optically pumped magnetometers or EEG to record responses to sensory perturbations across different anatomical areas, thus allowing one to estimate the sensory perturbational complexity index (28).

### Potential clinical impact

While we did not examine highrisk fetuses or infants, our results may nonetheless have important implications for developing predictive biomarkers of neuropsychiatric risk in early development. For example, prior work has shown that both spontaneous (1) and auditory-evoked (oddball) (57) EEG signal complexity in infants is predictive of later autism spectrum disorder (ASD) diagnoses. Given that earlier interventions generally lead to better outcomes for children with autism phenotypes (67), it may be especially beneficial to detect risk for ASD and other childhood psychiatric disorders (e.g., attention deficity hyperactivity disorder or ADHD) as early as the first weeks of life so that early preventative therapies can be administered before phenotypes emerge. Additionally, in cases of preterm birth, which are known to confer risk for neuropsychiatric disorders (16, 18), early neural markers of treatment responses based on signal complexity (21, 22) may be useful for improving therapies. All of these efforts may benefit from our investigation of neural complexity in response to sensory perturbations during the perinatal period, which lays the groundwork for future efforts by first establishing how neural complexity evolves very early in typical development.

Although it is still unknown precisely how neural entropy during the perinatal period may relate to neuropsychiatric risk, optimal healthy functioning of neural circuits may rest in a central ‘sweet spot’ or critical point which allows for both functional specialization and flexibility when entropy is neither too high nor too low. Very high entropy, as seen in spontaneous neural signals recorded from adults during the psychedelic state (10, 68) may indicate circuit flexibility and adaptability, but perhaps also an endophenotype associated with impaired Bayesian learning (69) that impedes healthy, adaptive behaviors, e.g., as seen in adults with schizophrenia (15, 70). Infants have not yet developed a strong Bayesian prior of the world (71), and so their neural activity is likely to be highly entropic (indeed, this may partially explain our finding of higher entropy in earlier development in the context of a Bayesian learning task). However, as the infant learns statistical regularities in the world, neural entropy should generally decrease in typical development as cortical circuits are sculpted by useful patterns of activity.

On the other hand, very low entropy may plausibly indicate that cortical circuit have become stuck in maladaptive activity patterns, corresponding, for instance, to repetitive or restricted behaviors see in obsessive compulsive disorder (OCD), ASD, ADHD, and Tourette syndrome (72–75). Both behavioral and neural activity patterns in these disorders can be usefully understood in the context of canalization (76): genetically programmed events and/or powerful experiences have lead the individual into a dysfunctional state delineated by the narrow contours of a metaphorical canal in neural and behavioral space. Notably, many of these disorders, such as ASD (77), OCD (78), ADHD (79), and Tourette syndrome (80) are diagnosed more frequently in boys. In light of this fact, and the overall greater sensitivity of males to risk factors in early development (59), it is reasonable to expect that boys and girls will, on average, show differences in neural entropy during childhood and even infancy. However, it remains unknown whether these factors indeed explain the sex finding in our data.

Although our sample of infants and fetuses was recruited from healthy mothers/expecting-mothers with normal-risk pregnancies, subjects still differed along relative risk factors whose correlations with neural variables may inform future efforts to derive risk biomarkers. Besides the higher relative risk incurred by male fetal sex (59), which corresponded to lower entropy (Table 1), mothers also varied with respect to the risk factor of BMI (53–55). Maternal BMI did correspond to lower synchronization in neonatal ERSPs (Fig. 3G), suggesting a less mature developmental profile in our data from infants born to relatively obese mothers. Similarly, a recent study by another fetal MEG site (34) found that visual and auditory evoked responses were significantly harder to detect in fetal MEG recorded from expecting mothers with a higher BMI; however, this could be explained by the presence of additional adipose tissue, which impedes stimulus transmission and may add distance between the fetal brain and the MEG sensors, making fetal brain activity harder to detect. We advocate for further work exploring the relationship between fetal MEG variables (e.g., entropy, evoked responses) and maternal risk factors for later psychiatric disorders in offspring.

Finally, while we studied healthy fetuses, recent work (25) suggests at least some convergence with data in preterm infants (a high risk population). Specifically, two spontaneous signals derived from frontal fNIRS data (the de-oxygenated hemoglobin and the cerebral tissue oxygenation index) also decrease in entropy as preterm infants mature (25). Thus, the entropy trajectory we found in sensory-evoked MEG from healthy fetuses may also occur, under certain contexts, in high risk populations such as preterm infants; however, the entropy of spontaneous EEG recorded from the same scalp site in these infants increased, rather than decreased, with PMA. As such, more work is required to understand which cortical signals grow or, alternatively, diminish in complexity in early development and why.

### Limitations, future directions, and conclusions

We have presented the first ever evidence suggesting that complexity in the cortical signals of human fetuses (measured as entropy) is linked to sex-specific developmental trajectories. We acknowledge several limitations of our study, including the relatively low SNR of our recordings and between-session variability in the head positioning of subjects. These limitations precluded more sophisticated analyses which would depend, e.g., on spatial information, such as functional connectivity or source localization (81), or on more permissive filtering, such as the broadband spectral exponent (48). Furthermore, we did not study preterm infants or other high risk samples of fetuses or infants. Lastly, we did not follow subjects past the neonatal period to track their later developmental and mental health phenotypes in early childhood, which would have helped us to better understand our results with respect to neuropsychiatric risk profiles.

We also acknowledge that even if the complexity of fetal and/or neonatal MEG signals offers a window onto sex-specific developmental trajectories, it may be challenging in practice to relate these data to later mental health outcomes due to the limitations of current technology (e.g., inflexible sensory placement, limited SNR). However, we expect that this may change in the near future, especially for newborns, due to advances in optically pumped magnetometers that will allow for optimal sensor placement and spatial coverage (28, 82).

A notable strength of our study was the inclusion of a large sample of fetuses with both cross-sectional and longitudinal sampling to study development, with longitudinal sampling continuing after birth in more than a third of the fetal subjects. Additionally, based on prior analyses of both our fetal and neonatal samples, we have strong reason to believe that our MEG data contain evoked responses reflecting cognitive processing and the early development of neural circuits that underpin hierarchical learning (30, 31).

In conclusion, our work builds on an earlier proof-of-concept study (56) to demonstrate the successful application of MEG entropy measures as markers of sex-specific developmental processes in fetuses. Future investigations should determine whether cortical signal entropy measured during the perinatal period is predictive of mental health outcomes in early childhood.

## Materials and Methods

### Study population

This work utilized publicly available data (see here for fetal data and here for neonatal data) which were acquired for and previously analyzed in studies of hierarchical learning in fetuses (31) and newborns (30) that revealed markers of perceptual consciousness in both cohorts. Previous studies from which the data were obtained had recruited parents or mothers-to-be who volunteered data from their newborn infants (N = 33) or fetuses (N = 60), respectively. Some mothers-to-be gave fetal data at multiple visits (see below). Newborns were 13 − 59 days old at the time of MEG recording. The studies were approved by the local ethics committee of the Medical Faculty of the University of Tübingen. Consent to participate in the experiment was signed by the mother-to-be (fetal group) or both parents (neonatal group).

### MEG recordings

Fetal and neonatal cortical signals were recorded in the context of earlier studies (see above) using the SARA (SQUID array for reproductive assessment, VSM MedTech Ltd., Port Coquitlam, Canada) system in a magnetically shielded room (Vakuumschmelze, Hanau, Germany) at the fMEG Center at the University of Tübingen. The SARA system is an MEG machine built for recording fetal data, with SQUID (superconducting quantum interference device) sensors arranged in a concavity that fits the maternal abdomen. All MEG signals were sampled at 610.3516 Hz; this legacy sampling rate originates from the need to avoid interference from other radio signals during system installation.

The SARA system is completely noninvasive and utilizes 156 primary sensors and 29 reference sensors. Because the sensor array, built to compliment the maternal abdomen, is much larger than the fetal or neonatal head, in both cases cortical signals are received largely by a subset of sensors that are located near the head. Based on simulation work (36) at the fMEG Center at the University of Tübingen, these optimally positioned sensors have an SNR of 9.2 − 11.6 dB after preprocessing and trial-averaging. Recorded signals included both cortical activity (MEG) and cardiac activity (MCG), the latter of which was retained to measure HRV and arousal. Fetal head position was determined before and after each MEG recording using ultrasound (Ultrasound Logiq 500MD, GE, UK), and the first measurement was used to place a positioning coil on the maternal abdomen. Three additional positioning coils were placed around the abdomen (left and right sides, and one on the spine) to track changes in the position of the mother-to-be’s body. Recordings with excessive positional changes were discarded. The SARA system was also used to record neonatal cortical signals by placing newborns in a crib resting head-first toward the sensor array. All newborns were positioned lying on their left side, such that auditory stimuli could be delivered to the right ear.

### Experiments

Experiments with fetuses and newborns used a local-global paradigm with pure tones (200 ms each) of two frequencies, 500 and 750 Hz. For each participant, one frequency was assigned as a standard tone and the other as a deviant tone. Assignments were kept consistent for participants with longitudinal visits, but varied randomly across different participants. Each experiment consisted of two blocks in randomized order, counter-balanced across participants: in one block, fetuses and newborns were trained with 30 sequences each consisting of either four identical standard tones (block rule: ‘ssss’) or three identical standard tones followed by a deviant (block rule: ‘sssd’). In each block, the tone duration was 200 ms, the inter-tone interval was 400 ms, and each sequence lasted 2000 ms, with a 1700 ms silent interval between sequences (1A). After this exposure phase, each block concluded with a testing phase of 180 sequences. In each test phase, 135 sequences (75%) were congruent with the block rule (global standard), whereas 45 sequences (25%) violated the block rule (global deviant). At a minimum, at least two global standards were included between each global deviant, and the order of sequences in each testing phase was otherwise pseudorandomized. Each block was approximately 13 minutes in duration. Because the fourth tone of each sequence can be compared to either other tones within the same sequence (local) or tones from the sequence introduced by the block rule (global), two levels of deviation are possible in this paradigm. For instance, given the block rule ‘sssd’, test phase sequence ‘sssd’ is a global standard but also a local deviant (i.e., the fourth tone is incongruent with the first three tones in its sequence—a first-order rule violation—but congruent with the global block rule sequence). Conversely, given the same block rule ‘sssd’, the test phase sequence ‘sssS’ (where the capital letter denotes a global rule violation) is a local standard but also a global deviant. For the block rule ‘ssss’, however, the test phase sequence ‘sssD’ is both a local and a global deviant. Given the hierarchical nature of the localglobal paradigm, it is possible that first and second-order rule violations produce different sizes of cortical perturbations. For additional details and protocols of this experiment, please see prior publications (30, 31).

### Data retention

Prior to preprocessing, MEG datasets were rejected from N = 4 participants in the fetal group and N = 6 participants in the neonatal group whose recordings were interrupted early. Of the remaining N = 56 participants in the fetal group, longitudinal recordings from N = 22 participants (N = 7 x two sessions, N = 3 x three sessions, and N = 12 x four sessions) were obtained. This yielded a total of 105 fetal datasets. Of these, fetal cortical signals were detected based on amplitude in 81 datasets which were retained (N = 24 x one session, N = 5 x two sessions, 9 x three sessions, N = 5 x four sessions) across 43 unique fetal subjects. Fetal cortical signals were identified based on a principal component approach and quality checked for consistency in location with fetal head position and overlap with remaining heart or muscle artifacts [see supplementary material of (31) for a detailed description of the process]. Of the fetal subjects retained, two were later born moderately preterm (GA > 32 weeks) and one very preterm (GA = 30 weeks), though this case of very preterm birth was due to the mother’s health rather than that of the fetus. These subjects were not studied after birth, as only healthy, full-term infants were recruited for the neonatal cohort.

All recordings from newborns were cross-sectional. As stated above, N = 6 newborns did not complete the full experiment, and thus data from N = 27 newborns entered preprocessing. After quality control, data from N = 20 newborns were retained, including 16 newborns who were also recorded from as fetuses.

### Data preprocessing

Although the input SNR of fetal MEG is extremely low (−60 to -20 dB), our previously validated preprocessing pipeline (31) yields enormous gains, resulting in a final SNR of 10.4 dB (average based on simulated signals) (36), which is comparable to the raw signal from high quality recordings of adult MEG (83). The SNR for neonatal MEG recordings is substantially higher than that of fetuses, as the infant’s head can be positioned immediately next to the sensory array, avoiding any impediment from the maternal abdomen or the intermixing of cortical signals with maternal heart signals. In both cases (fetal and neonatal), prior published work (30, 31) has already demonstrated event-related signatures of hierarchical learning using the same data and preprocessing that we utilized in the current project. These pre-processing pipelines are summarized below, though we also direct the interested reader to prior publications for full details (30, 31, 36, 84).

### MEG signal preprocessing

Preprocessing of MEG data was conducted using MATLAB R2016b (The MathWorks, Natick, MA, USA). Using a 4th-order Butterworth filter, fetal signals were bandpass filtered at 1-10 Hz and neonatal signals were bandpass filtered at 1-15 Hz prior, which is a typical filtering range for fetal and neonatal event related responses (85). For each fetal recording, a cluster of 10 channels with the highest signal amplitude after artifact removal were root mean square averaged, normalized as a percentage change from the first auditory evoked response, and chosen for further analysis. Similarly, for each neonatal recording, five channels containing cortical signals were identified using principal component analysis, room mean square averaged, and chosen for further analysis (86). Signals were segmented from -200 ms to +3000 ms referenced to the onset of the first tone (Fig. S7). To adjust for the excess number of standard trials in each block, only standard trials immediately preceding deviant trials were analyzed. Note that we did not examine shorter subsegments of trial-averaged signals (e.g., after the onset of the fourth tone) due to the lowpass filtering which limits the usefulness of shorter data segments (e.g., in fetal data lowpass filtered at 10 Hz, a 1000 ms subsegment would contain at most 10 oscillatory cycles).

### Heart rate variability (HRV)

As a proxy for arousal level, HRV was measured in all fetal and neonatal data. R peaks were detected in the fetal/neonatal MCG signal prior with the FLORA algorithm (84) prior to MCG signal isolation using the FAUNA algorithm (36). The R peaks were then used to compute the normal-to-normal R-R intervals (87), the standard deviation of which (SDNN) was taken as our measure of HRV. SDNN was log10 transformed prior to statistical analysis to better approximate a Gaussian distribution.

### Signal entropy

Data analysis from this point forward was conducted using MATLAB R2021b/2022a. Entropy is an information-theoretic quantity which measures the degree of uncertainty (or unpredictability) in data generated by a given probability distribution. When applied to time series data, it can be interpreted as quantifying the degree of diversity in a signal: the higher a signal’s entropy, the more it will explore different trajectories. It is also, in a sense, a measure of information, as highly entropic signals may have a greater informational content (88). To estimate MEG signal entropy, we used several approaches applied to signals that had already been averaged across trials and channels from each recording. First, we utilized the LZC (37) and CTW (38) compression algorithms, which determine the number of unique substrings in the signal, a quantity which relates to the signal’s ground truth entropy (more technically, to its entropy rate) (88). Both approaches require the signal to be transformed into discrete symbols. For both LZC and CTW, we satisfied this requirement in a binary fashion by thresholding each signal using its median value. CTW returns the entropy rate of the signal estimated from a variable-order Markov Model.

Besides compression-based methods, we also utilized entropy estimates based on state-space reconstruction (89). These include the modified sample entropy (mSampEn) (39) and permutation entropy (PermEn) (41). The mSampEn reflects the tendency of motifs to reoccur within a signal (90) within a certain tolerance or state space radius r. Our calculations of mSampEn used *r* = 0.15*σ*, where *σ* is the standard deviation of the signal. We also examined a variant of mSampEn, the modified multiscale sample entropy, or mMSE, which computes mSampEn at different timescales using a coarse graining procedure (40); specifically, we averaged mMSE across 20 timescales. At each timescale, we updated r based on the coarse grained signal [for a rationale, see (91, 92)]. Both mSampEn and mMSE were calculated using z-scored signals. The PermEn reflects the occurrence of unique permutations based on ordinal rankings of data (41), a distinction with important implications for the results of our entropy decomposition, as small changes in the signal which affect the ordinal rankings of data points may have large effects as measured by PermEn. These ordinal rankings are created by choosing a lag or timescale to separate samples. We computed PermEn using two different lags (93, 94) appropriate for our data: *τ* = 32 ms (PermEn32, sensitive to 4 - 10 Hz activity) and *τ* = 64 ms (PermEn64, sensitive to 2 - 5 Hz activity) and an embedding dimension of m = 3. We then normalized PermEn by the quantity log(m!) such that *PermEn* ∈ [0, 1]. All entropy measures were computed after averaging MEG channels of interest (see “Data retention” above).

### Cortical dynamics

To investigate whether the dynamics of each cortical signals were stochastic (i.e., containing intrinsic randomness), or deterministic (i.e., predetermined by initial conditions, encompassing both periodic and chaotic behavior), we adapted a decision tree algorithm by Toker et al (33). The algorithm infers whether signal dynamics are stochastic using two rounds of surrogate data testing with 1000 surrogates. For the first round, 1000 surrogates were generated using the iterative amplitude adjusted Fourier Transform method that exactly preserves the power spectrum (IAAFT-2). Additionally, to rule out the possibility of nonlinear stochasticity, a second round of surrogate data testing was performed using 1000 cyclic phase permutation surrogates. Because de-noising is crucial to the algorithm’s success (33), we lowpass filtered signals from newborns again at the same frequency (10 Hz) as had been applied to fetal signals (4th-order Butterworth filter) before classifying their dynamics. Having identified all signals as either stochastic or deterministic, LMMs were then used to predict signal dynamics (see “Statistical analysis” below).

### Statistical modeling of signal entropy and dynamics

To model signal entropy, we selected predictors using stepwise regression with the MATLAB function stepwiselm using default parameters. For each entropy measure, the model started with a constant term (intercept), and the following variables were made available as predictors for both fetuses and newborns using the stepwise procedure: age (gestational age if the subject was a fetus), HRV (measured as SDNN), sex, birth weight, gestational age at birth, maternal age, maternal BMI (measured before pregnancy), block rule, and stimulus. A final common model for all entropy measures was then constructed separately for fetuses and newborns using all predictor terms which were selected for at least half of the entropy measures (i.e., if a predictor term was present in at least three out of six entropy models, it was included in the final common model). The final common model was constructed as a LMM using the MATLAB function fitlme, thus accounting for multiple stimuli, block rules, and laboratory visits for each subject using random intercepts.

To test *K* predictors of entropy measured *N* times across *n* subjects, LMMs used the formula

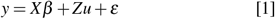

where *y* is the *N* x 1 vector of entropy estimates for each measure, *X* is the *N* x *K* matrix of fixed effects, including a fixed intercept term (first column), *β* is the *K* x 1 vector of fixed effect coefficients, *Z* is the *N* x *n* design matrix of random effects, *u* is the *n* x 1 vector of random intercepts, and *ε* is the *N* x 1 vector of residuals.

After running the stepwise procedure, the final common LMM for fetal subjects consisted of the model intercept plus following fixed effects: gestational age, sex, maternal age, and the interaction of gestational age with sex. For neonatal subjects, the final common LLM consisted of the model intercept plus age as a fixed effect term.

Having chosen separate models for fetal and neonatal entropy, we next examined entropy measures derived from neonatal signals that were lowpass filtered at the same frequency (10 Hz) as fetal signals. This allowed us to pool neonatal and fetal signals together and select models for the pooled data using a stepwise procedure. When entering age into models, we used the postmenstrual age (i.e., gestational age for fetuses and postbirth age plus gestatioanl age at birth for newborns). Each stepwise procedure began with a base model that included an intercept plus a categorical variable indicating whether the subject was a fetus or a newborn. In this case, our final LMM including the above terms plus fixed effects of postmenstrual age and sex, as well as random effects (intercepts) for each subject.

Next, to evaluate the effect of surrogacy on signal complexity measures (see “Surrogate data testing” below), we used LMMs to model entropy by including the same fixed effects as above as covariates, plus a fixed effect of surrogacy (treated as a binary variable indicating whether signals are surrogates).

Additionally, to model signal dynamics, we used LMMs with the same fixed effects as used in the final common model for entropy measures, but with dynamics (i.e., a categorical variable indicating the outcome of the decision tree for categorizing signal dynamics) as the dependent variable.

### Statistical modeling of signal power

To compute ERSPs that map cortical responses to tones in frequency and time, we used Morlet wavelets with 8 wavelets per octave, yielding a total of 28 wavelets logarithmically spaced from 1 to 10.5 Hz (fetal data) and 32 wavelets logarthimically spaced from 1 to 15 Hz (neonatal data), with 44 timepoints linearly spaced from -200 to 3000 ms (referenced to the time of the first tone). To analyze ERSPs, we modeled log_10_(power) at each element of the TFR using predictors selected with lasso regression. Predictors considered by the lasso regression were age/GA for newborns/fetuses, HRV, sex, MA, PMA at birth, birth weight, maternal BMI, block rule, and stimulus. A regularization parameter *λ* was used to penalize model complexity, and interaction terms were not considered as predictor variables. The lasso regression model consolidated all TFR elements into a single response vector. First, we explored a parameter space of 25 *λ* values logarthmically spaced between 0.02 and 2 (fetuses) and 0.033 and 1 (newborns); note that we explored a different parameter range for newborns as the values used for fetuses did not sufficiently reduce model complexity in newborns. Next, the *λ* value which yielded optimal model performance as judged by 10-fold cross validation was selected. We then fit a new lasso regression to the response vector using the chosen *λ* value. Predictor variables with non-zero regularization coefficients were then selected and used in LMMs with random intercepts to subsequently model spectral power at each TFR element. This procedure was performed separately for fetal and neonatal data. For fetuses, the lasso regression yielded GA and stimulus as predictor variable entered as a fixed effect into LMMs. For newborns, the lasso regression yielded age, PMA at birth, MA, SDNN, and maternal BMI as predictor variables entered as fixed effects into LMMs.

Here, we faced a multiple comparisons problem as models were fit for multiple time-frequency elements. To ovrcome this issue we used threshold free cluster enhancement (TFCE) with parameters recommended by Mensen and Khatami (95), E = 2/3 and H = 2. This approach favors time-frequency elements that are surrounded by neighboring elements displaying similar effects. Moreover, unlike previous cluster statistical methods (96), TFCE considers many different thresholds instead of a single arbitrary threshold. In our case, we sampled 25 evenly spaced thresholds for test statistics starting from 0 and increasing to the largest test statistic present in the data (as judged by absolute value). We then used permutation testing to derive an empirical P-value for each electrode. Data were permuted by randomly shuffling the outcome variable, i.e., log_10_(power), for each of 2000 permutations. For all thresholds explored, elements were considered neighbors if their edges or corners were touching.

### Entropy decomposition

We used an entropy decomposition (44) to investigate the extent to which amplitude versus non-amplitude (i.e., phase and its interaction with amplitude) properties of the MEG signal contributed to changes in each entropy measure. The decomposition requires two sets of paired data to compute differences in entropy. For each signal, we generated 250 surrogates (97) by shuffling phases separately within younger versus and subjects in one procedure and then between younger and older subjects pooled together in another procedure. By applying the inverse Fourier transform to obtain surrogate signals from these shuffling procedures, the decomposition algorithm allows for the effects of amplitude, phase, and their interaction to be systematically disentangled, e.g., when phases are pooled between age groups, the difference that remains is due to amplitude. For full details of the algorithm, please refer to Mediano et al. (43).

In fetal MEG recordings, we performed the decomposition using only those fetuses with longitudinal data from both relatively early and late timepoints, based on the lower (< 32 weeks) and upper (> 35 weeks) tertiles (i.e., thirds) of GA. Accordingly, we identified n = 12 fetuses with data from both tertiles delineated above. After decomposing entropy changes into amplitude, phase, and amplitude x phase interaction components, we summed the differences in each entropy measure attributable to phase and phase x amplitude interactions to create a single “non-amplitude” quantity Data from all four conditions (i.e., global rule x stimulus combinations) were used, and we modeled the entropy differences Δ*H* using LMMs with a fixed effect of condition and random intercepts for subjects; we then added a fixed effect of the signal property (amplitude or non-amplitude) and used a log-likelihood ratio test to evaluate whether adding this term significantly improved the model fit.

We also applied the entropy decomposition on neonatal data; however, due to the smaller sample size of newborns, we performed a median split on the data (rather than a tertile split), grouping all newborns that were recorded from at 34 days of age and younger separately from those recorded from at older ages; this enabled us to performed the decomposition in a larger number of newborns (n = 10 in each subgroup) than if we had only sampled the lower and upper tertiles, though at the expense of maintaining an age buffer in between the two subgroups. Furthermore, because each subject was only recorded from at most once as a newborn, we were unable to perform the neonatal entropy decomposition within-subjects. As a workaround, for each of the older newborns, we computed the average difference between its entropy and that of the younger newborns; this procedure was then repeated with each of the younger newborns compared to older newborns. In other words, for the *ith* newborn in the younger subgroup, we computed an entropy difference Δ*H* between its signal and the signal of all older newborns according to

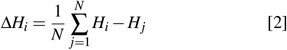

where *j* indexes each of the older newborns. Similarly, for the *jth* newborn in the older subgroup, we computed Δ*H* between its signal and the signal of all younger newborns according to

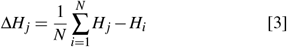

and so we obtained one average difference for each participant, allowing us to ask which component of the entropy change was larger for each participant; note that the sum of Δ*H*_*i*_ across all *N* newborns in the younger subgroup equals the sum of Δ*H*_*j*_ across all *N* newborns in the older subgroup, i.e., the referencing of entropy differences to the younger or older subgroup is equivalent and therefore arbitrary.

Following the above procedure, we computed the average difference between the entropy of each neonatal recording in one subgroup and the entropy of the recordings in the other age group. Next, we evaluated whether meaningful differences existed between amplitude and non-amplitude contributions using the same log-likelihood ratio test approach described above for fetal data.

### Mediation analysis

To determine whether the effect of sex on fetal entropy was mediated by mean signal amplitude, we ran a separate nonparametric mediation analysis in R (version 3.6.2) for each fetal entropy measure using the Mediation package (44) with 1000 bootstrapped resamples. Each analysis compared LMMs [implemented using the lme4 package(98)] with and without the mediator variable.

### Surrogate data testing

To assess whether the entropy of cortical signals differed from that of noise with similar spectral properties (i.e., surrogate data), we utilized surrogate data testing (97, 99). Surrogate signals were generated from each cortical signal using the iterative amplitude adjusted Fourier transform that exactly preserves the amplitude distribution (IAAFT-1) algorithm (100), resulting in signals with maximal entropy given a fixed variance (70). Note that we followed best practices by generating the surrogate signal after truncation of the original signal, so that start and end points had approximately the same value and same first derivative (99). The entropy of cortical signals was calculated separately with truncation only for the purpose of comparing with surrogates (i.e., elsewhere, we used the entropy computed from the full signal without truncation). For each cortical signal, entropy measures were also computed for each of 100 surrogate signals; the median entropy was then computed across all surrogates. To evaluate whether a given measure differed between cortical and surrogate signals, we used LMMs (see “Statistical analysis” above) and evaluated the surrogacy term in each model.

### Correlations between MEG measures

To investigate possible relationships amongst MEG measures, we examined correlations between signal entropy measures both with each other and with spectral power. For neonatal data, correlations were computed using Pearson correlation coefficients. Because fetal data show statistical dependencies between longitudinal recordings (see Table S4), we z-scored MEG measures from fetal data and used the normalized beta coefficients from random intercept regression models. Because standardized betas in LMMs depend on the variance of the random effect and are thus generally asymmetrical (i.e., *β*_*i, j*_ ≠*β* _*i, j*_), we used the mean of *β* _*i, j*_and *β* _*i, j*_ to represent the correlation between entropy measure i and j (Fig. S1). This was done using LMMs which predicted a first measure *M*_*i*_ from a second measure *M* _*j*_ (fixed effect) and random intercepts, and vice versa (reversing the roles of *M*_*i*_ and *M* _*j*_), where *M*_*i*_ and *M* _*j*_ are the pair of MEG measures whose correlations is being determined. We then averaged the beta coefficients *β* _*i, j*_ and *β* _*i, j*_ corresponding to *M*_*i*_ and *M* _*j*_ to estimate the correlation strength. Note that *β* ∈ [−1, 1] because *M*_*i*_ and *M* _*j*_ have unit variance after z-scoring.

Because our concern was mostly with correlation direction and size rather than statistical significance, we did not derive p-values for correlations. Besides investigating correlations across recordings, we also examined temporal correlations of CTW with spectral power after averaging the ERF across all datasets within each of four stimulus/rule conditions. A notable property of CTW is its superior temporal resolution over LZC due to its faster convergence, which can be used to evaluate sub-second changes in entropy (45). For this reason, we used time-resolved CTW with 164 ms (i.e., 100 sample) sliding windows with 90% overlap to evaluate correlations between entropy and spectral power on the grand-averaged ERFs. Spectral power was computed using Morlet wavelet to yield the same temporal resolution as CTW, and in each case the frequency resolution was set to four wavelets per octave (1 - 10 Hz, fetal data; 1 - 15 Hz, neonatal data).

### Spectral exponent

We also investigated correlations between entropy measures and the slope of the aperiodic component of the MEG spectrum, i.e., the spectral exponent. The spectral exponent was measured using the fitting oscillations and one-over-f (FOOOF) algorithm (101). Considering our lowpass filter choices, we modeled the 1 - 9.5 Hz spectrum for fetuses and the 1 - 14.5 Hz spectrum for newborns, with the spectrum obtained by time-averaging the TFRs derived from Morlet wavelets as described previously. In the FOOOF settings, we set peak limits to 0.2 - 2.5 Hz and we set the aperiodic mode to ‘fixed’. As above, we did not derive p-values for correlations because our concern was mostly with correlation direction and size rather than statistical significance and, moreover, because we could not obtain broadband estimates of the spectral exponent due to earlier signal filtering. Correlations shown in Fig. S3G are Pearson coefficients obtained by treating repeated measures as independent samples (we did not observe substantial differences between Pearson coefficients and beta coefficients in the correlation matrix of fetal entropy measures in Fig. S4C and therefore opted for Pearson correlations as the more intuitive approach).

## ACKNOWLEDGMENTS

We are grateful to all volunteers and families who participated in our research. We would also like to thank Daniel Toker for his input on methodological aspects of our study, Alessandra DallaVecchia for her input on “sensory PCI”, Simon Ruch for assisting with a code review, and Tim Bayne for many enlightening discussions on infant consciousness. Finally, we thank the fMEG team for their contributions, including Franziska Schleger (original study design), and Magdalene Weiss (data collection). Additionally, we gratefully acknowledge the following funders: 1) the German Federal Ministry of Education and Research (BMBF) grants Somnia (13GW0294), Enable (13GW0359) and Bevares (13GW0570), 2) the European Union’s Joint Programme for Neurodegenerative Disease Research (EU-JPND 2022-130) grant Recast (01ED2309) 3) the FET Open Luminous project (H2020 FETOPEN-2014-2015-RIA under agreement No. 686764) as part of the European Union’s Horizon 2020 research and 2014 – 2018 training program, 4) the German Federal Ministry of Education and Research (BMBF) to the German Center for Diabetes Research (DZD01GI0925), 5) the Deutsche Forschungsgemeinschaft (DFG, German Research Foundation; 493345456), 6) the Wellcome Trust (grant no. 210920/Z/18/Z), and 7) the Open Access Publishing Fund of the University of Tübingen.

## License

This work is licensed under a [Creative Commons Attribution 4.0 International License](https://creativecommons.org/licenses/by/4.0/).

## References

1. WJ Bosl, H Tager-Flusberg, CA Nelson, Eeg analytics for early detection of autism spectrum disorder: a data-driven approach. Sci. reports 8, 1–20 (2018).

2. C Gu, ZX Liu, S Woltering, Electroencephalography complexity in resting and task states in adults with attention-deficit/hyperactivity disorder. Brain Commun. 4, fcac054 (2022).

3. S Guan, et al., The complexity of spontaneous brain activity changes in schizophrenia, bipolar disorder, and adhd was examined using different variations of entropy. Hum. Brain Mapp. 44, 94–118 (2023).

4. M Aljalal, SA Aldosari, M Molinas, K AlSharabi, FA Alturki, Detection of parkinson’s disease from eeg signals using discrete wavelet transform, different entropy measures, and machine learning techniques. Sci. Reports 12, 22547 (2022).

5. D Abásolo, et al., Analysis of regularity in the eeg background activity of alzheimer’s disease patients with approximate entropy. Clin. neurophysiology 116, 1826–1834 (2005).

6. AG Casali, et al., A theoretically based index of consciousness independent of sensory processing and behavior. Sci. translational medicine 5, 198ra105–198ra105 (2013).

7. J Frohlich, et al., Neural complexity is a common denominator of human consciousness across diverse regimes of cortical dynamics. Commun. Biol. 5, 1374 (2022).

8. S Sarasso, et al., Consciousness and complexity during unresponsiveness induced by propofol, xenon, and ketamine. Curr. Biol. 25, 3099–3105 (2015).

9. M Dasilva, et al., Modulation of cortical slow oscillations and complexity across anesthesia levels. Neuroimage 224, 117415 (2021).

10. C Timmermann, et al., Human brain effects of dmt assessed via eeg-fmri. Proc. Natl. Acad. Sci. 120, e2218949120 (2023).

11. FX Vollenweider, KH Preller, Psychedelic drugs: neurobiology and potential for treatment of psychiatric disorders. Nat. Rev. Neurosci. 21, 611–624 (2020).

12. D Toker, et al., Consciousness is supported by near-critical slow cortical electrodynamics. Proc. Natl. Acad. Sci. 119, e2024455119 (2022).

13. PC Ivanov, et al., Multifractality in human heartbeat dynamics. Nature 399, 461–465 (1999).

14. CS Poon, CK Merrill, Decrease of cardiac chaos in congestive heart failure. Nature 389, 492–495 (1997).

15. T Takahashi, et al., Antipsychotics reverse abnormal eeg complexity in drug-naive schizophre-nia: a multiscale entropy analysis. Neuroimage 51, 173–182 (2010).

16. S Johnson, et al., Psychiatric disorders in extremely preterm children: longitudinal finding at age 11 years in the epicure study. J. Am. Acad. Child & Adolesc. Psychiatry 49, 453–463 (2010).

17. S Johnson, N Marlow, Preterm birth and childhood psychiatric disorders. Pediatr. Research 69, 11–18 (2011).

18. C Nosarti, et al., Preterm birth and psychiatric disorders in young adult life. Arch. general psychiatry 69, 610–617 (2012).

19. S Janjarasjitt, M Scher, K Loparo, Nonlinear dynamical analysis of the neonatal eeg time series: the relationship between sleep state and complexity. Clin. neurophysiology 119, 1812–1823 (2008).

20. MS Scher, H Waisanen, K Loparo, MW Johnson, Prediction of neonatal state and maturational change using dimensional analysis. J. clinical neurophysiology 22, 159–165 (2005).

21. F Kaffashi, M Scher, S Ludington-Hoe, K Loparo, An analysis of the kangaroo care intervention using neonatal eeg complexity: a preliminary study. Clin. neurophysiology 124, 238–246 (2013).

22. JR Isler, RI Stark, PG Grieve, MG Welch, MM Myers, Integrated information in the eeg of preterm infants increases with family nurture intervention, age, and conscious state. PloS One 13, e0206237 (2018).

23. AB Barrett, AK Seth, Practical measures of integrated information for time-series data. PLoS computational biology 7, e1001052 (2011).

24. O De Wel, et al., Complexity analysis of neonatal eeg using multiscale entropy: applications in brain maturation and sleep stage classification. Entropy 19, 516 (2017).

25. L Semeia, et al., Multiscale entropy analysis of combined eeg-fnirs measurement in preterm neonates. bioRxiv pp. 2023–07 (2023).

26. C Sortica da Costa, et al., Complexity of brain signals is associated with outcome in preterm infants. J. Cereb. Blood Flow & Metab. 37, 3368–3379 (2017).

27. RL Goldenberg, JF Culhane, JD Iams, R Romero, Epidemiology and causes of preterm birth. The lancet 371, 75–84 (2008).

28. J Frohlich, et al., Not with a “zap” but with a “beep”: measuring the origins of perinatal experience: Origins of perinatal experience. NeuroImage p. 120057 (2023).

29. V Rajagopalan, et al., Local tissue growth patterns underlying normal fetal human brain gyrification quantified in utero. J. neuroscience 31, 2878–2887 (2011).

30. J Moser, et al., Magnetoencephalographic signatures of hierarchical rule learning in newborns. Dev. cognitive neuroscience 46, 100871 (2020).

31. J Moser, et al., Magnetoencephalographic signatures of conscious processing before birth. Dev. cognitive neuroscience 49, 100964 (2021).

32. G Tononi, M Boly, M Massimini, C Koch, Integrated information theory: from consciousness to its physical substrate. Nat. Rev. Neurosci. 17, 450–461 (2016).

33. D Toker, FT Sommer, M D’Esposito, A simple method for detecting chaos in nature. Commun. biology 3, 11 (2020).

34. H Eswaran, et al., Tracking evoked responses to auditory and visual stimuli in fetuses exposed to maternal high-risk conditions. Dev. psychobiology 63, 5–15 (2021).

35. L Semeia, K Sippel, J Moser, H Preissl, Evaluation of parameters for fetal behavioural state classification. Sci. reports 12, 3410 (2022).

36. K Sippel, et al., Fully automated subtraction of heart activity for fetal magnetoencephalography data in 2019 41st Annual International Conference of the IEEE Engineering in Medicine and Biology Society (EMBC). (IEEE), pp. 5685–5689 (2019).

37. A Lempel, J Ziv, On the complexity of finite sequences. IEEE Transactions on information theory 22, 75–81 (1976).

38. FM Willems, YM Shtarkov, TJ Tjalkens, The context-tree weighting method: Basic properties. IEEE transactions on information theory 41, 653–664 (1995).

39. HB Xie, WX He, H Liu, Measuring time series regularity using nonlinear similarity-based sample entropy. Phys. Lett. A 372, 7140–7146 (2008).

40. M Costa, AL Goldberger, CK Peng, Multiscale entropy analysis of complex physiologic time series. Phys. review letters 89, 068102 (2002).

41. C Bandt, B Pompe, Permutation entropy: a natural complexity measure for time series. Phys. review letters 88, 174102 (2002).

42. SM Smith, TE Nichols, Threshold-free cluster enhancement: addressing problems of smoothing, threshold dependence and localisation in cluster inference. Neuroimage 44, 83–98 (2009).

43. PA Mediano, FE Rosas, AB Barrett, D Bor, Decomposing spectral and phasic differences in nonlinear features between datasets. Phys. Rev. Lett. 127, 124101 (2021).

44. D Tingley, T Yamamoto, K Hirose, L Keele, K Imai, Mediation: R package for causal mediation analysis. (2014).

45. Y Gao, I Kontoyiannis, E Bienenstock, Estimating the entropy of binary time series: Methodology, some theory and a simulation study. Entropy 10, 71–99 (2008).

46. R Gao, EJ Peterson, B Voytek, Inferring synaptic excitation/inhibition balance from field potentials. Neuroimage 158, 70–78 (2017).

47. JD Lendner, et al., An electrophysiological marker of arousal level in humans. Elife 9, e55092 (2020).

48. N Schaworonkow, B Voytek, Longitudinal changes in aperiodic and periodic activity in electrophysiological recordings in the first seven months of life. Dev. cognitive neuroscience 47, 100895 (2021).

49. LC Shuffrey, et al., Aperiodic electrophysiological activity in preterm infants is linked to subsequent autism risk. Dev. Psychobiol. 64, e22271 (2022).

50. PJ Marshall, Y Bar-Haim, NA Fox, Development of the eeg from 5 months to 4 years of age. Clin. neurophysiology 113, 1199–1208 (2002).

51. J Huo, SF Quan, J Roveda, A Li, Coupling analysis of heart rate variability and cortical arousal using a deep learning algorithm. Plos one 18, e0284167 (2023).

52. CB Saper, PM Fuller, NP Pedersen, J Lu, TE Scammell, Sleep state switching. Neuron 68, 1023–1042 (2010).

53. HM Rivera, KJ Christiansen, E. Sullivan, The role of maternal obesity in the risk of neuropsychiatric disorders. Front. neuroscience p. 194 (2015).

54. AG Edlow, Maternal obesity and neurodevelopmental and psychiatric disorders in offspring. Prenat. diagnosis 37, 95–110 (2017).

55. F Cirulli, C Musillo, A Berry, Maternal obesity as a risk factor for brain development and mental health in the offspring. Neuroscience 447, 122–135 (2020).

56. J Moser, et al., Evaluating complexity of fetal meg signals: a comparison of different metrics and their applicability. Front. systems neuroscience 13, 23 (2019).

57. FC Peck, et al., Prediction of autism spectrum disorder diagnosis using nonlinear measures of language-related eeg at 6 and 12 months. J. neurodevelopmental disorders 13, 1–13 (2021).

58. E Hisle-Gorman, et al., Prenatal, perinatal, and neonatal risk factors of autism spectrum disorder. Pediatr. research 84, 190–198 (2018).

59. JA DiPietro, KM Voegtline, The gestational foundation of sex differences in development and vulnerability. Neuroscience 342, 4–20 (2017).

60. MD Wheelock, et al., Sex differences in functional connectivity during fetal brain development. Dev. cognitive neuroscience 36, 100632 (2019).

61. KM Cook, et al., Robust sex differences in functional brain connectivity are present in utero. Cereb. Cortex 33, 2441–2454 (2023).

62. S Sarasso, et al., Consciousness and complexity: a consilience of evidence. Neurosci. Conscious. 7, 1–24 (2021).

63. S Lippé, N Kovacevic, R McIntosh, Differential maturation of brain signal complexity in the human auditory and visual system. Front. human neuroscience p. 48 (2009).

64. AR McIntosh, N Kovacevic, RJ Itier, Increased brain signal variability accompanies lower behavioral variability in development. PLoS computational biology 4, e1000106 (2008).

65. B Mišic, T Mills, MJ Taylor, AR McIntosh, Brain noise is task dependent and region specific. J. Neurophysiol. 104, 2667–2676 (2010).

66. GN Elston, T Oga, I Fujita, Spinogenesis and pruning scales across functional hierarchies. J. Neurosci. 29, 3271–3275 (2009).

67. EB Itzchak, D. Zachor, Who benefits from early intervention in autism spectrum disorders? Res. Autism Spectr. Disord. 5, 345–350 (2011).

68. MM Schartner, RL Carhart-Harris, AB Barrett, AK Seth, SD Muthukumaraswamy, Increased spontaneous meg signal diversity for psychoactive doses of ketamine, lsd and psilocybin. Sci. reports 7, 46421 (2017).

69. H Rajpal, et al., Psychedelics and schizophrenia: Distinct alterations to bayesian inference. NeuroImage 263, 119624 (2022).

70. D Bai, W Yao, S Wang, J Wang, Multiscale weighted permutation entropy analysis of schizophrenia magnetoencephalograms. Entropy 24, 314 (2022).

71. A Gopnik, The philosophical baby: What children’s minds tell us about truth, love & the meaning of life. (Random House), (2009).

72. M Jiujias, E Kelley, L Hall, Restricted, repetitive behaviors in autism spectrum disorder and obsessive–compulsive disorder: A comparative review. Child Psychiatry & Hum. Dev. 48, 944–959 (2017).

73. C Ganos, T Ogrzal, A Schnitzler, A Münchau, The pathophysiology of echopraxia/echolalia: relevance to gilles de la tourette syndrome. Mov. Disord. 27, 1222–1229 (2012).

74. D Efron, RC Dale, Tics and tourette syndrome. J. paediatrics child health 54, 1148–1153 (2018).

75. A Mogadam, et al., Magnetoencephalographic (meg) brain activity during a mental flexibility task suggests some shared neurobiology in children with neurodevelopmental disorders. J. Neurodev. Disord. 11, 1–12 (2019).

76. R Carhart-Harris, et al., Canalization and plasticity in psychopathology. Neuropharmacology p. 109398 (2022).

77. E Fombonne, Epidemiological surveys of autism and other pervasive developmental disorders: an update. J. autism developmental disorders 33, 365–382 (2003).

78. DJ Castle, A Deale, IM Marks, Gender differences in obsessive compulsive disorder. Aust. & New Zealand J. Psychiatry 29, 114–117 (1995).

79. CU Greven, JS Richards, JK Buitelaar, Sex differences in adhd. Oxf. textbook attention deficit hyperactivity disorder pp. 154–160 (2018).

80. J Garris, M Quigg, The female tourette patient: sex differences in tourette disorder. Neurosci. & Biobehav. Rev. 129, 261–268 (2021).

81. P Nevalainen, et al., Somatosensory evoked magnetic fields from the primary and secondary somatosensory cortices in healthy newborns. Neuroimage 40, 738–745 (2008).

82. MJ Brookes, et al., Magnetoencephalography with optically pumped magnetometers (opm-meg): the next generation of functional neuroimaging. Trends Neurosci. (2022).

83. A Jaiswal, et al., Comparison of beamformer implementations for meg source localization. NeuroImage 216, 116797 (2020).

84. K Sippel, et al., Fully automated r-peak detection algorithm (flora) for fetal magnetoencephalographic data. Comput. methods programs biomedicine 173, 35–41 (2019).

85. F Schleger, et al., Magnetoencephalographic signatures of numerosity discrimination in fetuses and neonates. Dev. neuropsychology 39, 316–329 (2014).

86. J Moser, K Sippel, F Schleger, H Preißl, Automated detection of fetal brain signals with principal component analysis in 2019 41st Annual International Conference of the IEEE Engineering in Medicine and Biology Society (EMBC). (IEEE), pp. 6549–6552 (2019).

87. H Mat Husin, et al., Maternal weight, weight gain, and metabolism are associated with changes in fetal heart rate and variability. Obesity 28, 114–121 (2020).

88. TM Cover, Elements of information theory. (John Wiley & Sons), (1999).

89. CJ Stam, Nonlinear dynamical analysis of eeg and meg: review of an emerging field. Clin. neurophysiology 116, 2266–2301 (2005).

90. JS Richman, JR Moorman, Physiological time-series analysis using approximate entropy and sample entropy. Am. journal physiology-heart circulatory physiology (2000).

91. VV Nikulin, T Brismar, Comment on “multiscale entropy analysis of complex physiologic time series”. Phys. review letters 92, 089803 (2004).

92. A Humeau-Heurtier, The multiscale entropy algorithm and its variants: A review. Entropy 17, 3110–3123 (2015).

93. JR King, et al., Information sharing in the brain indexes consciousness in noncommunicative patients. Curr. Biol. 23, 1914–1919 (2013).

94. P Bourdillon, et al., Brain-scale cortico-cortical functional connectivity in the delta-theta band is a robust signature of conscious states: an intracranial and scalp eeg study. Sci. reports 10, 14037 (2020).

95. A Mensen, R Khatami, Advanced eeg analysis using threshold-free cluster-enhancement and non-parametric statistics. Neuroimage 67, 111–118 (2013).

96. E Maris, R Oostenveld, Nonparametric statistical testing of eeg-and meg-data. J. neuroscience methods 164, 177–190 (2007).

97. J Theiler, S Eubank, A Longtin, B Galdrikian, JD Farmer, Testing for nonlinearity in time series: the method of surrogate data. Phys. D: Nonlinear Phenom. 58, 77–94 (1992).

98. D Bates, et al., Linear mixed-effects models using eigen and s4. r package version 1.1-23 (2013).

99. G Lancaster, D Iatsenko, A Pidde, V Ticcinelli, A Stefanovska, Surrogate data for hypothesis testing of physical systems. Phys. Reports 748, 1–60 (2018).

100. T Schreiber, A Schmitz, Improved surrogate data for nonlinearity tests. Phys. review letters 77, 635 (1996).

101. T Donoghue, et al., Parameterizing neural power spectra into periodic and aperiodic components. Nat. neuroscience 23, 1655–1665 (2020).

