## Supplemental Material for "Sex differences in prenatal development of neural complexity in the human brain"

**This PDF file includes:**

Figs. S1 to S4
Tables S1 to S4

### 1. Supplemental Figures

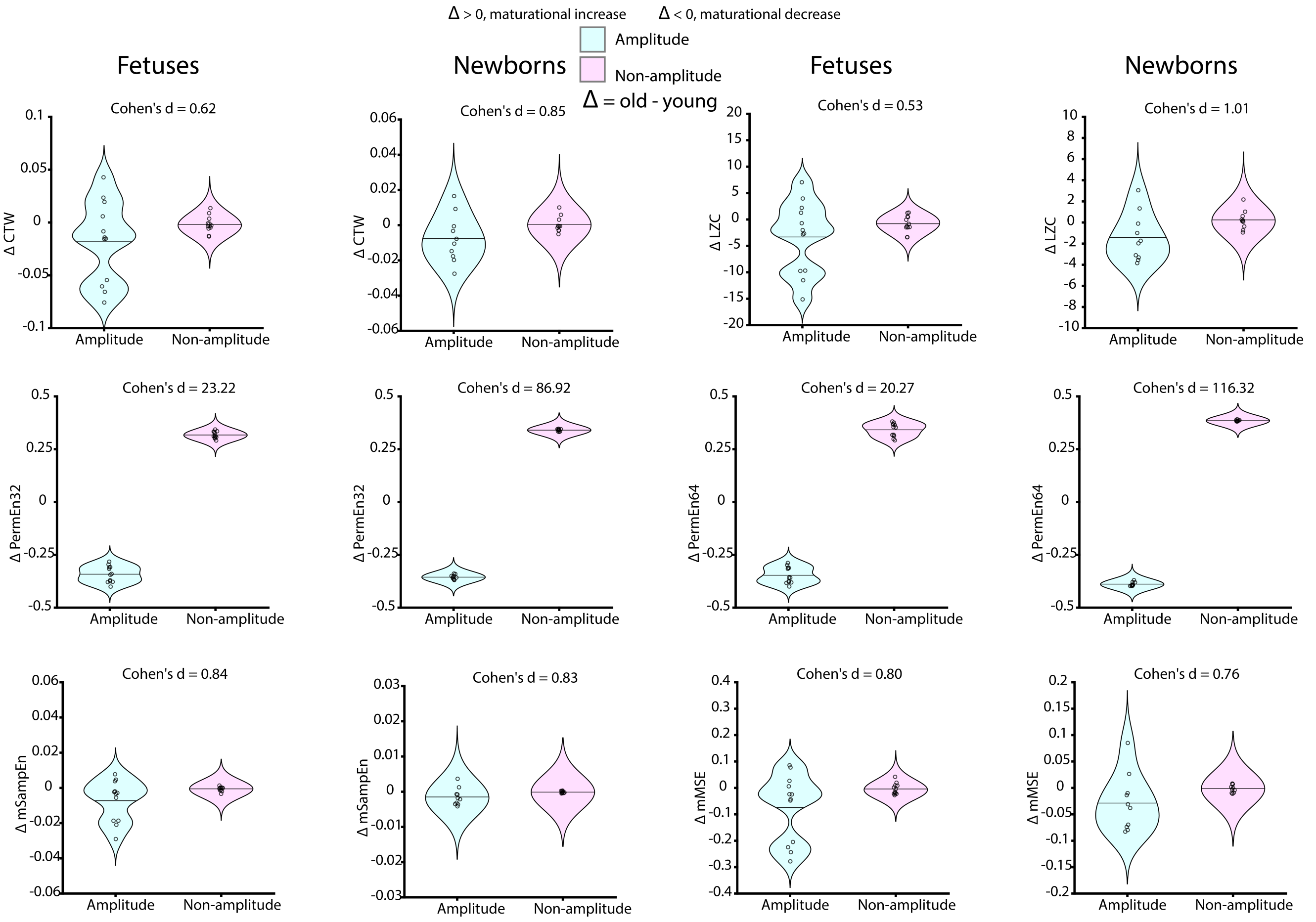


**Supplemental Figure 1: Changes in entropy resulting from amplitude or non-amplitude signal changes.** Data distributions in each panel are averaged across all four experimental conditions (i.e., block rule and stimulus combinations). PermEn32 and PermEn64 (middle row) showed a far larger dissociation between amplitude and non-amplitude signal properties, likely because the PermEn algorithm is sensitive to small changes in the signal which result in new ordinal rankings of data points. Effect sizes (Cohen's d) are indicated for each decomposition.

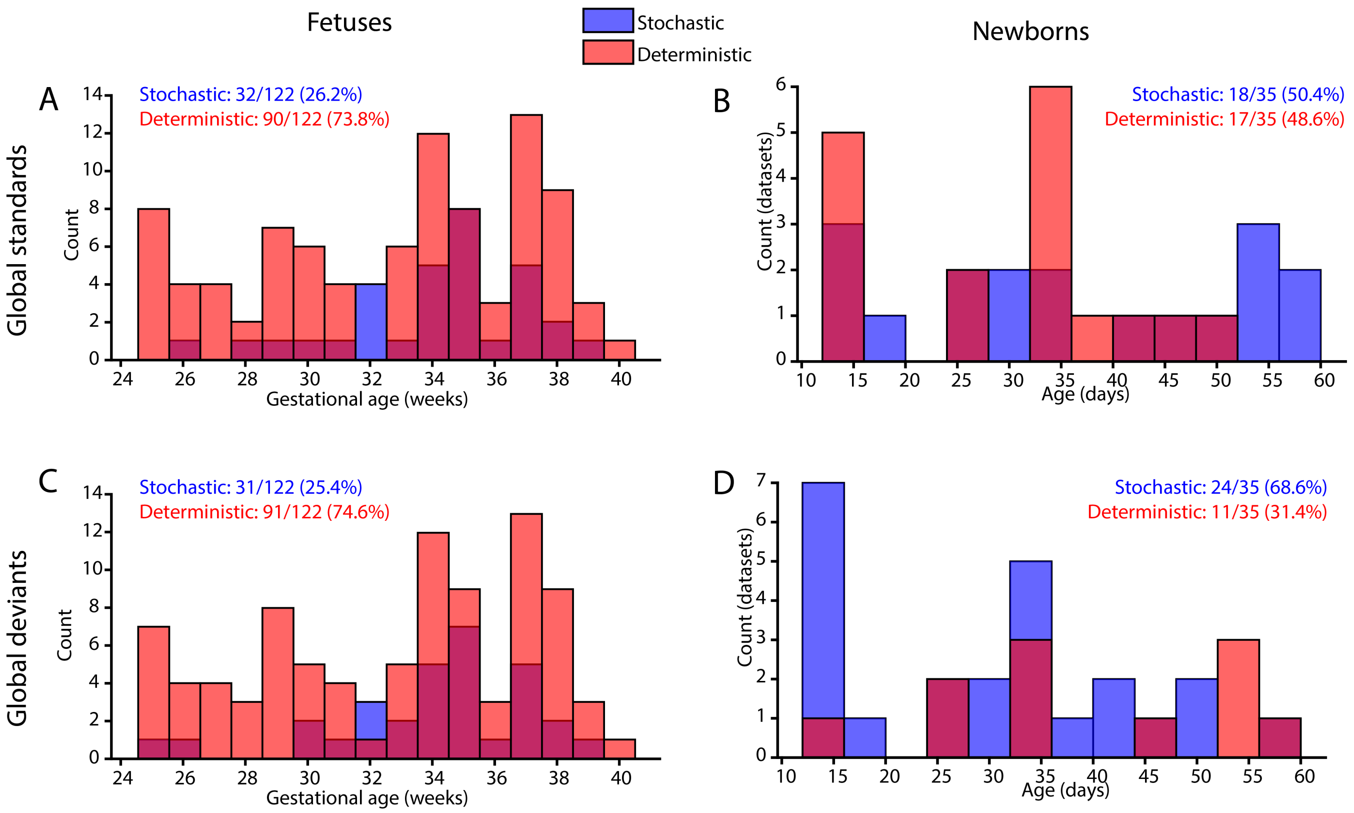


**Supplemental Figure 2: Histograms of signal dynamics categories (stochastic or deterministic) by gestational age (fetuses, left column) and age (newborns, right column).** The first two rows show results from global standards (A, B) and the second row shows results from global deviants (C, D). Both fetuses and newborns displayed a mixture of stochastic and deterministic dynamics. In fetuses, the majority of recordings were deterministic, whereas in newborns, the majority of recordings were stochastic. Dynamics were not significantly predicted by maturation in either group, though the proportion of recordings with stochastic dynamics was significantly higher in newborns than in fetuses (chi-squared test, $\chi^2 = 28.6$, $P = 9.1 \cdot 10^{-8}$).

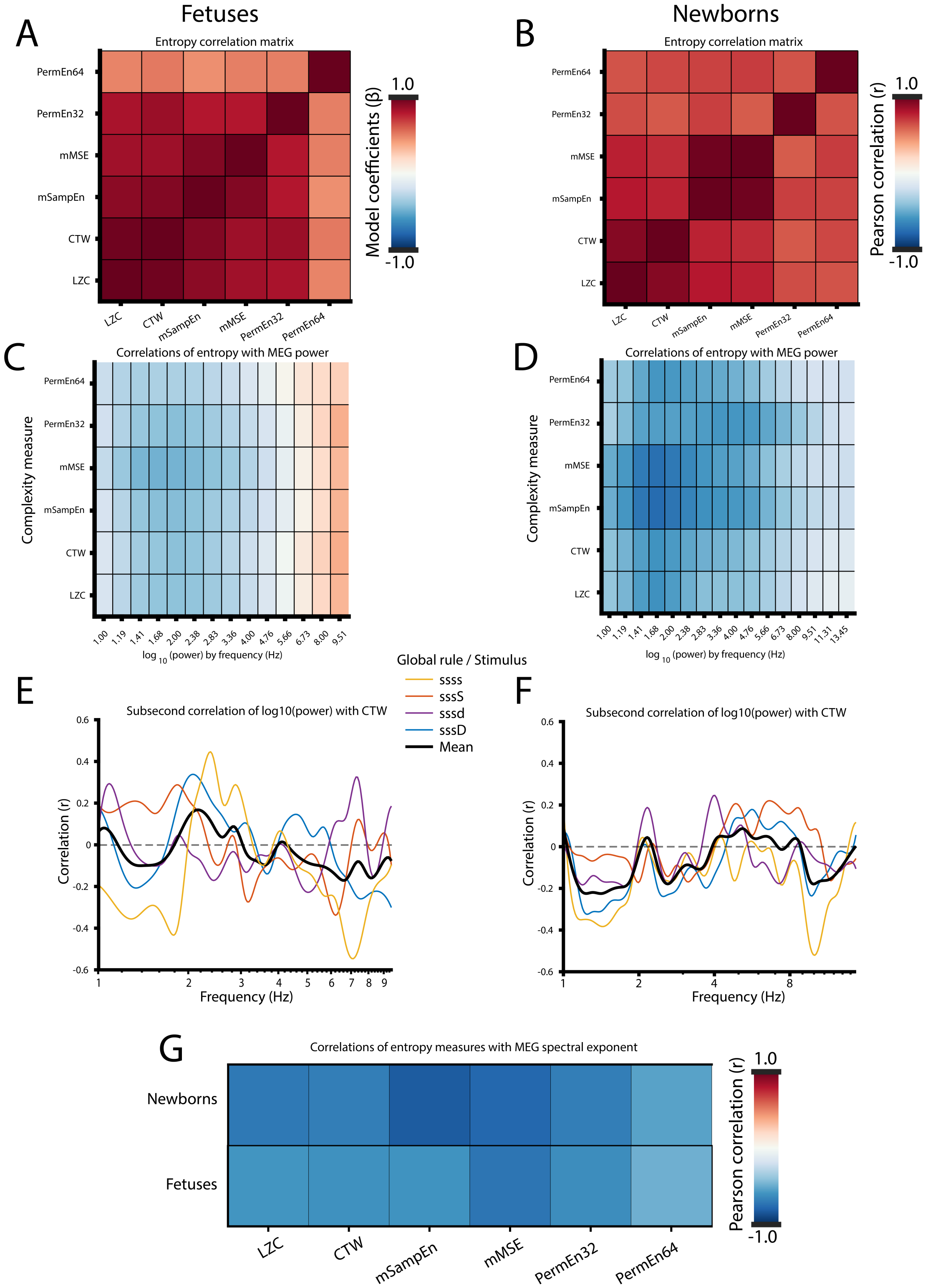


**Supplemental Figure 3: Correlations between MEG measures.** Entropy measures were highly correlated with one another in both fetuses (A) and newborns (B). These same entropy measures show negative correlations with spectral power at most frequencies in fetuses (C) and all frequencies in newborns (D). Subsecond CTW did not correlate strongly with subsecond spectral power after averaging across conditions in fetuses (E) or newborns (F). Correlations between entropy masures and the MEG spectral exponent were uniformerly negative (G).

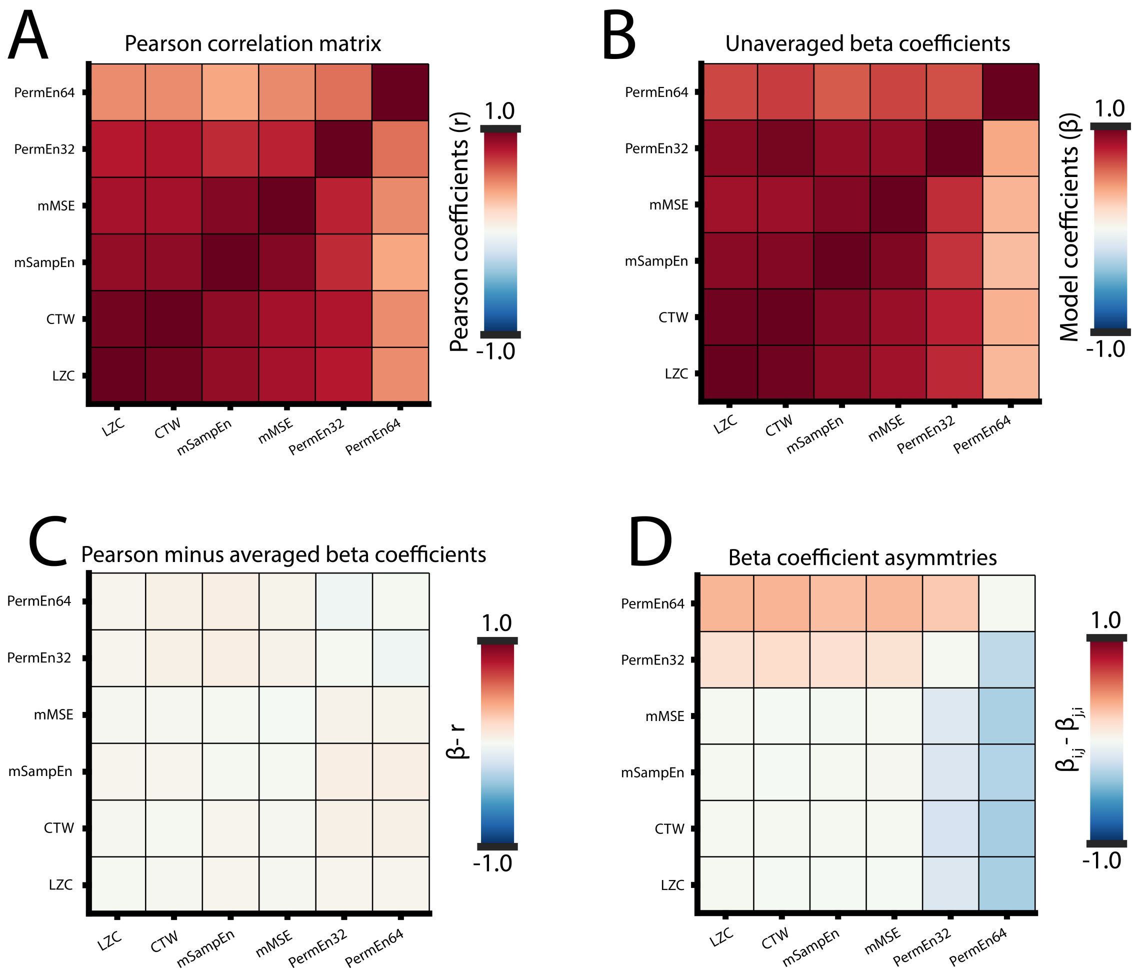


**Supplemental Figure 4: Alternative calculations of correlations in fetal entropy measusres.** Because our fetal data contained multiple recordings from the same fetal subjects and, moreover, the random effect term significantly increased the fits of most model predicting entropy, we did not wish to rely on Pearson correlation coefficients (A) between entropy measures from each recording, as these correlation estimates may overrepresent subjects with multiple recordings. For this reason, we instead utilized standardized model coefficients (betas) from linear mixed models that predicted entropy measures from each other while accounting for random effects (B). Differences between Pearson coefficients and averaged beta coefficients (C) are very small ($\beta - r < 0.1$ in all cases). However, standardized betas in LMMs are not generally symmetrical (i.e., $\beta - i, j \neq \beta_{j,i}$), since they depend on the variance of the random effect, and the random effect may often contribute more to one variable or the other. To address this problem, we used the mean of $\beta_{i,j}$ and $\beta_{j,i}$ to represent the correlation between entropy measure i and j. Here, for transparency, we show the asymmetry in the standardized betas prior to averaging (D), which primarily affected correlation estimates for PermEn64.

### 2. Supplemental Tables

#### Supplemental Table 1.

***Mean signal amplitude does not significantly mediate the effect of sex on entropy.***

| Measure | P-value |
|---|---|
| CTW | 0.61 |
| LZC | 0.75 |
| PermEn32 | 0.26 |
| PermEn64 | 0.48 |
| mMSE | 0.87 |
| mSampEn | 0.71 |

**Table S1. Mean signal amplitude does not significantly mediate the effect of sex on entropy for any fetal entropy measure (P > 0.05 all measures, even without accounting for multiple testing). We ran a separate analysis for each fetal entropy measure with 1000 Monte Carlo draws (nonparametric bootstraping) in each analysis.**

#### Supplemental Table 2.

***Surrogate data testing of MEG complexity measures in fetuses and newborns.***

| Statistic | CTW | LZC | SE | MSE | PermEn32 | PermEn64 |
|---|---|---|---|---|---|---|
| Fetus P-value | 0.0789 | 0.105 | 0.0555 | 0.434 | 0.0319 | 0.821 |
| Fetus t-stat | -1.76 | -1.62 | 1.92 | -0.782 | -2.15 | -0.227 |
| Fetus Survive FDR | FALSE | FALSE | FALSE | FALSE | FALSE | FALSE |
| Newborn P-value | 0.337 | 0.663 | 1 | 1 | 0.946 | 0.0154 |
| Newborn t-stat | -0.963 | -0.436 | 0 | 2.09e-13 | -0.0674 | 2.45 |
| Newborn survive FDR | FALSE | FALSE | FALSE | FALSE | FALSE | TRUE |

**Table S2. Entropy measures computed from cortical signals were compared to the median entropy across 100 surrogate signals. Significant differences between entropy from cortical and surrogates signals were assessed using linear mixed models (LMMs). Surrogacy only significantly predicted entropy using mSampEn in newborns ($P < P_{crit}$), but note also trend-level effects in fetuses for PermEn32 ($P < 0.05$), as well as mSampEn and CTW ($P < 0.1$)**

#### Supplemental Table 3.

***Predictors of MEG signal dynamics in fetuses and newborns.***

| Predictor | P-value |
|---|---|
| Fetal GA tstat | 1.87 |
| Fetal GA Pvalue | 0.0633 |
| Fetal GA survive FDR | FALSE |
| Fetal MA Pvalue | 0.0579 |
| Fetal MA tstat | -1.91 |
| Fetal Sex x GA Pvalue | 0.151 |
| Fetal Sex x GA tstat | -1.44 |
| Fetal Sex x GA survive FDR | FALSE |
| Fetal Sex Pvalue | 0.189 |
| Fetal Sex tstat | 1.32 |
| Fetal Sex survive FDR | FALSE |
| Neonatal age tstat | -0.218 |
| Neonatal age Pvalue | 0.828 |
| Neonatal age survive FDR | FALSE |

**Table S3. Signal dynamics (i.e., stochastic versus deterministic behavior) were not significantly predicted by any variable. Note, however, trend-level effects ($P < 0.1$) for GA and MA in fetuses.**

### Supplemental Table 4.

***Log-likelihood ratio tests between models with and without random intercepts for fetal subjects.***

| Measure | Log-liklihood ratio stat | P-value |
|---|---|---|
| CTW | 17.9 | 2.3e-05 |
| LZC | 16.6 | 4.67e-05 |
| PermEn32 | 18 | 2.17e-05 |
| PermEn64 | 0 | 1 |
| mMSE | 20 | 7.85e-06 |
| mSampEn | 17.6 | 2.73e-05 |

**Table S4. Prediction of all entropy measures except for PE64 is significantly improved by including random effects, thus demonstrating that longitudinal recordings from fetal subjects show statistical dependencies within-subject.**
